# Imp, a key regulator of transposable elements, cell growth, and differentiation genes during embryogenesis

**DOI:** 10.1101/2025.09.30.679481

**Authors:** Paula Vazquez-Pianzola, Dirk Beuchle, Peter M’Angale, Gimena Alegre, Greco Hernández, Simon L. Bullock, Travis Thomson, Beat Suter

**Affiliations:** Institute of Cell Biology, University of Bern, 3012 Bern, Switzerland; Department of Neurobiology, University of Massachusetts Chan Medical School, Worcester, Massachusetts, United States of America; mRNA and Cancer Laboratory, Instituto Nacional de Cancerología (INCan; National Institute of Cancer). Mexico City 14080, Mexico; Tecnologico de Monterrey, Escuela de Medicina y Ciencias de la Salud. Mexico; Division of Cell Biology, MRC Laboratory of Molecular Biology, Cambridge, UK

## Abstract

Imps are a highly conserved family of RNA-binding proteins involved in embryonic development, cancer progression, and neurogenesis. However, the molecular pathways and RNAs regulated by Imp to control these processes remain poorly understood. Embryos derived from *Imp* mutant germline clones arrest development, and transcriptome analysis revealed significant dysregulation of genes involved in cell growth, differentiation, tube morphogenesis, neuronal projection development, and RNA metabolism, along with de-repression of transposable element (TE) RNAs. Consistent with these findings, *Imp* mutant embryos display TE-overexpression phenotypes, are smaller in size, and exhibit defective organ development, including impaired tracheal branching and gastrulation. Reduced levels of Imp at the larval neuromuscular junction (NMJ) impair synaptic bouton formation and decrease adult longevity. RIP-seq experiments showed that Imp-associated RNAs are enriched for TE RNAs. Proteomic analyses confirmed that several TE-encoded proteins are upregulated in *Imp* mutant embryos. Specifically, the Ty1 family retrotransposon *Copia* was derepressed. Consistent with recent findings that *Copia* is a potent inhibitor of synaptogenesis, its upregulation likely contributes to the impaired NMJ formation and broader embryonic defects observed in *Imp* mutants. Moreover, Imp associates with piRNA pathway proteins, ensures Piwi nuclear localization, and—like *piwi* mutants—its loss disrupts TE silencing and causes position-effect variegation (PEV) defects. The analysis of Imp complexes further points to potential mechanisms by which Imp may regulate TE expression. Overall, these results indicate that Imp maintains genome stability and ensures proper developmental progression and neuronal activity by regulating post-transcriptional processes and suppressing transposons.

## INTRODUCTION

Imps (<u>I</u>GF2 <u>m</u>RNA-binding <u>p</u>roteins), originally named for their ability to bind insulin-like growth factor 2 (IGF2) mRNA, are a family of highly conserved RNA-binding proteins known to regulate a broad set of transcripts involved in development and disease. Roles in RNA localization, translation, and stability of specific RNAs have been described for the different Imp orthologs (Degrauwe et al., 2016; Vazquez-Pianzola and Suter, 2012). Mammals express three Imp paralogs (Imp1-3) and *Drosophila* codes for only one *Imp* gene, which is more closely related to Imp1 (Vazquez-Pianzola and Suter, 2012). The expression pattern of the mammalian Imp1 and Imp3 paralogs is oncofetal, whereas Imp2 expression persists in some normal adult tissues. Imp1 and Imp3 are also overexpressed in several cancer types (Degrauwe et al., 2016; Mancarella and Scotlandi, 2020). Like mammalian orthologs, *Drosophila Imp* is expressed throughout embryogenesis and has also been implicated in cancer progression, where it is required to sustain neuroblast tumor growth (Narbonne-Reveau et al., 2016; Nielsen et al., 2000).

The embryonic expression of Imp protein paralogs correlates with their essential role during embryogenesis (Xu et al., 2022). In zebrafish, depletion of maternal Imp3 destabilizes maternal mRNAs before the maternal-to-zygotic transition (MZT), leading to abnormal cytoskeletal organization, cell cycle defects, and impaired embryonic development(Ren et al., 2020). Conversely, overexpression of Imp3 inhibits MZT, resulting in developmental delay (Ren et al., 2020). In mice, depletion of Imp2 also causes early embryonic arrest by targeting CCAR1 and RPS14, and by stabilizing IGF2 mRNA during zygotic genome activation (ZGA) (Liu et al., 2019; Xu et al., 2022). A similar effect on IGF2 regulation has been reported in human embryos. Loss of *Imp1* in mice causes dwarfism and impaired gut development, with only 50% of the *Imp1* mutant progeny reaching the third day of life (Hansen et al., 2004). Mutant mice are 40% smaller than the wild type, with most organs hypoplastic. Vg1 RBP, the *Xenopus* ortholog, is essential for cell migration in early neural development and for maintaining epithelial integrity (Shoshkes et al., 2015; Yaniv et al., 2003). *Drosophila Imp* is also an essential gene during embryogenesis (Boylan et al., 2008; Munro et al., 2006). Females lacking maternal *Imp* exhibit normal ovarian development; however, they produce non-viable embryos, with the majority dying during early development and only a few surviving until late larval or early pupal stages (Boylan et al., 2008; Munro et al., 2006).

Imp protein orthologs have essential roles in RNA metabolism, principally in the transport and local translation of specific RNAs. *Xenopus* Imp (also referred to as Vg1RBP or Vera) binds *Vg1* mRNA and localizes with it to the vegetal pole of *Xenopus* oocytes during maturation (Deshler et al., 1998; Havin et al., 1998). Vg1RBP and chicken Imp1, also known as zipcode-binding protein 1 (ZBP-1), are required for the localization and translational repression of β-actin mRNA during its transport to the leading edge of motile fibroblasts and neurons (Hüttelmaier et al., 2005; Oleynikov and Singer, 2003; Yaniv et al., 2003). Proper localization of β-actin mRNA supports axon guidance, dendritic branching, and growth in developing neurons (Buxbaum et al., 2014; Lepelletier et al., 2017; Leung et al., 2006; Sasaki et al., 2010). In *Drosophila* brains, Imp assembles into neuronal RNP granules that include *Profilin* mRNA, which encodes an actin regulator (Medioni et al., 2014). The prion-like domain (PLD) of Imp, a motif associated with the neurodegenerative progression of other neuronal RNA-binding proteins, is essential for the developmentally regulated localization of Imp RNP granules to axons and for Imp’s role in axon remodeling (Vijayakumar et al., 2019). *Drosophila* Imp also binds RNAs critical for embryonic patterning, such as *osk* and *grk*. However, although these sequences are essential for the localization of these RNAs in oocytes and embryos, their localization and translation are unaffected in *Imp* mutant ovaries and early embryos (Boylan et al., 2008; Munro et al., 2006). These and other mRNAs essential for oocyte determination, differentiation, establishment of anterior-posterior, dorsal-ventral and apical-basal polarity are transported by a general RNA transport machinery in *Drosophila.* This machinery includes Bicaudal-D (BicD) and the RNA-binding protein Egalitarian (Egl), which interact with cytoplasmic microtubule motors—Dynein and Dynactin—to move mRNA cargo along microtubules to distinct cellular compartments (Bullock and Ish-Horowicz, 2001; Dienstbier et al., 2009; Vazquez-Pianzola et al., 2017).

The role of Imp, if any, within the BicD/Egl/Dynein localization machinery has not been investigated. Moreover, the molecular mechanisms underlying the essential maternal function of *Imp* in embryonic development, as well as the regulatory networks in which Imp operates, remain unclear. Imp RNA targets have been systematically identified in larval brains, cultured cells, and in other biological systems (Hansen et al., 2004; Hansen et al., 2015; Lee et al., 2025). However, the embryonic RNA targets regulated by *Imp* in *Drosophila*, and particularly those that could explain the developmental arrest observed in embryos lacking maternal Imp, have yet to be comprehensively defined.

In this study, we identify RNAs bound by Imp and characterize the transcriptomic and proteomic differences between wild-type and *Imp* mutant embryos. We show that loss of maternal *Imp* leads to widespread gene dysregulation, affecting genes involved in growth, differentiation, morphogenesis, and RNA metabolism. Notably, *Imp* mutant embryos exhibit strong de-repression of transposable element (TE) RNAs and proteins, particularly those of the Ty1/Copia retrotransposon family, which are known to impair synaptogenesis. These embryos are smaller in size, display defective gastrulation and tracheal development. Furthermore, larvae with reduced Imp levels at the neuromuscular junction (NMJ) show impaired synaptic bouton formation and a reduced adult lifespan. All these phenotypes are consistent with known consequences of TE misregulation. Mechanistically, Imp binds TE transcripts, interacts with components of the piRNA pathway, and is required for the nuclear localization of Piwi. Furthermore, *Imp* mutants exhibit defects in position-effect variegation (PEV), linking Imp to the regulation of heterochromatin and transposon silencing. Together, these findings suggest that Imp safeguards genome integrity and supports normal developmental progression by coordinating post-transcriptional regulation and transposon repression.

## RESULTS

### *Drosophila* Imp makes a complex with the BicD/dynein transport machinery

Previous mass spectrometry analyses identified Imp in BicD immunoprecipitates (Vazquez-Pianzola et al., 2011, reproduced in Fig. S1A). The presence of Imp in BicD/Egl complexes was further confirmed by reverse immunoprecipitation experiments (Fig. 1A). Antibodies targeting GFP co-immunoprecipitated Imp-GFP along with BicD, Egl and poly-A binding protein (PABP), in Imp-GFP^G080^ embryo extracts expressing a functional Imp-GFP trap protein (Fig. 1A, Fig. S2). Imp was also specifically pulled down using anti-GFP antibodies from extracts expressing Egl::GFP (Fig. 1A). These findings show that Imp is a constituent of the BicD/Egl complex. Antibodies targeting BicD co-immunoprecipitated Imp, Egl and PABP. Treatment with RNase reduced the BicD/Imp interaction by 84 % (Fig. S1B, Vazquez-Pianzola et al., 2011). The Egl/BicD interaction was not affected, as expected for a known direct interaction between these two proteins (Fig. S1B, (Dienstbier et al., 2009; Vazquez-Pianzola et al., 2011). This suggests that the presence of Imp in the BicD-RNP is at least in part dependent on or stabilized by the associated RNAs. To test for additional direct interaction between BicD and Imp, we performed a yeast two-hybrid assay. We observed a weak but consistent direct interaction between the carboxy terminus of BicD and Imp proteins (Fig. 1B).

**Fig. 1.**
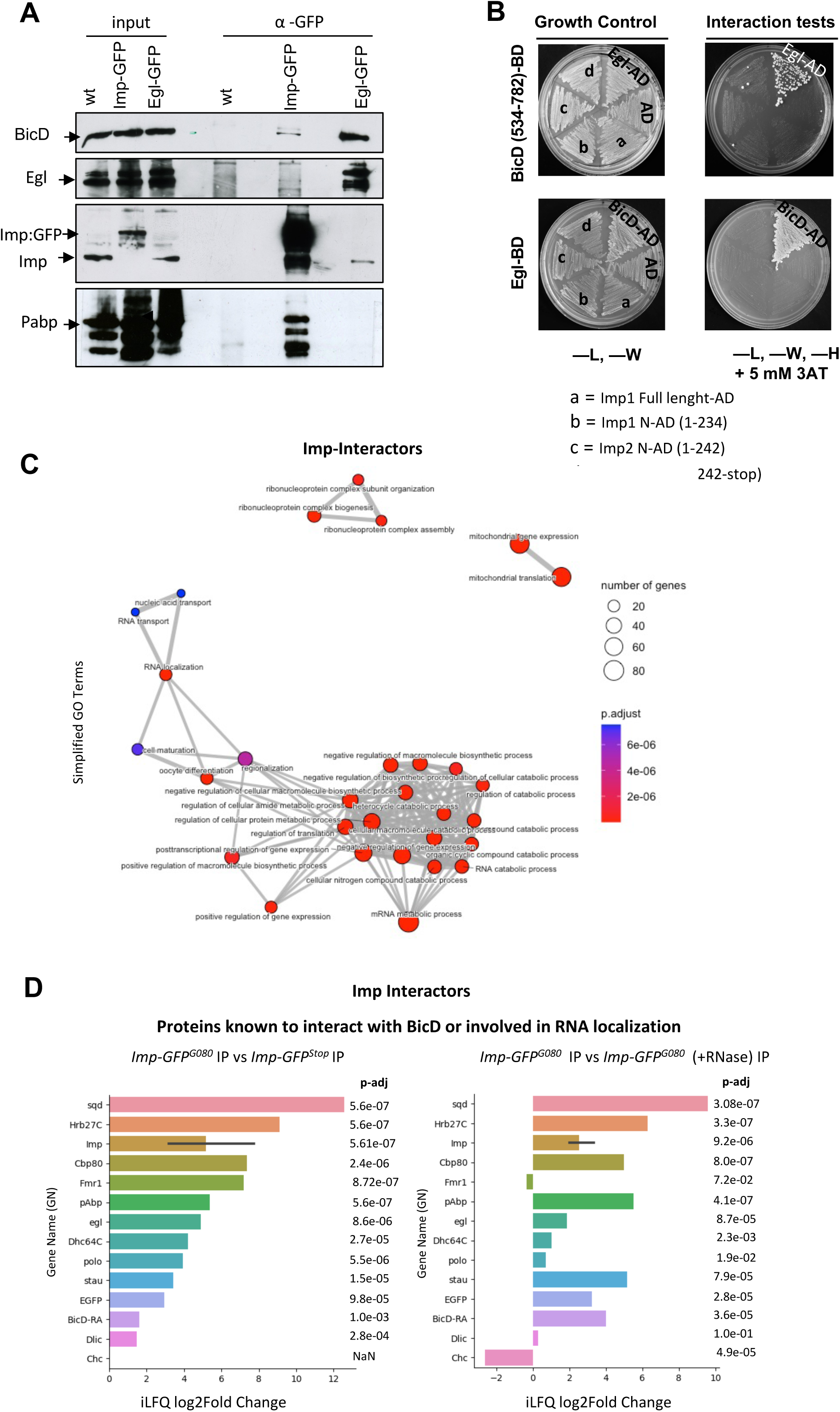
Imp forms part of the BicD/Egl complex and GO analysis of Imp interactors shows association with proteins involved in RNA metabolic pathways. **(A)** Immunoprecipitation (IP) of wild-type (wt, used as mock IP control), Imp-GFP^G080^, and Egl-GFP expressing embryos with anti-GFP antibodies. The IP material was tested for the presence of BicD, Egl, Imp, and Pabp. 0.15% of the cytoplasmic extract used for each IP was loaded as an input control. **(B)** Imp weakly interacts with BicD but not with Egl in a yeast two-hybrid assay. Interaction test of BicD carboxy terminus (top, amino acids 534-782) and full-length Egl (lower) with different fragments of Imps in the activator domain (AD). *Drosophila Imp* produces 2 isoforms differing in the first N-terminal amino acids (18 or 22 different amino acids at the N-term), described here as Imp1 and Imp2. Imp1 full-length (a), Imp1 amino terminus (amino acids 1-234) (b), Imp2 amino terminus (amino acids 1-242) (c), and IMP1/2 carboxy-terminal (amino acids 235-566, shared in both isoforms) (d) were tested. Egalitarian-AD and BicD full-length-AD were used as positive controls, respectively, and empty vector AD as a negative control. A weak but consistent interaction between the carboxy-terminal part of BicD and Imp1/2 carboxy-terminal was detected. **(C)** Embryo extracts from *Imp-GFP^G080^*, *Imp-GFP^Stop^* or *yw* were subjected to immunoprecipitation (IP) using anti-GFP antibodies. Additionally, *Imp-GFP^G080^* extracts were subjected to IP with anti-GFP antibodies in the presence of RNase to detect direct interactors. The IP materials were directly analyzed by bulk mass spectrometry. GO analysis of the proteins found enriched in the *Imp-GFP^G080^*IP compared to the mock IPs (performed with *yw* and *Imp-GFP^Stop^* extracts). Proteins were selected based on a log2Fold Change >1 or were uniquely present in the *Imp-GFP ^G080^,* IP, with a padj <0.05. Simplified GO Terms identified with Cluster Profiler are shown. **(D)** Enrichment of selected proteins known to be involved in RNA transport in oogenesis and proteins known to be in complex with BicD and in Imp complexes are shown. The log2FoldChange enrichment between the peptides found in the *Imp-GFP^G080^* IP and the mock control IP (*Imp-GFP^Stop^*) and between the peptides found in the *Imp-GFP^G080^*IP done in the absence and presence of RNase is shown together with the corresponding padj values of the enrichment.

### Identification of novel Imp interactors

To identify novel proteins that associate with Imp that could give us an insight into the essential roles of maternal IMP during embryo development, we conducted immunoprecipitation (IP) experiments using embryo extracts expressing the Imp-GFP^G080^ trap protein followed by mass spectrometry analysis (Morin et al., 2001). As a mock control for the immunoprecipitation experiments, we generated a mutant version of the functional *Imp-GFP*^G080^ chromosome, using a CRISPR-Cas9 mutagenesis screen. The selected mutant chromosome has an 8-nucleotide deletion after the GFP insertion site, leading to a frameshift in the open reading and the introduction of a premature stop codon (Fig. S2, see methods). This results in the production of the fusion protein Imp*^1-31-^*GFP-Imp^32-95-^- XX^18aa^- Stop protein expressed under the same regulatory domains as the *Imp-GFP*^G080^ line (Fig. S2B). This mutant allele is lethal in flies. We refer to this fly line as *Imp-GFP^Stop^*and use it as a mock control for the immunoprecipitations (Fig. S2A,B). Thus, extracts from embryos expressing either Imp-GFP^G080^, Imp-GFP^Stop^, or wild-type embryos were subjected to immunoprecipitation (IP) with anti-GFP antibodies followed by mass spectrometry analysis of the immunoprecipitated material (Fig. S3, raw data in Tables S1-2).

The IPs were performed in the presence and absence of RNase A to identify factors that bind independently of the presence of RNA in the complex. Fig. S3 shows the top 100 interactors, and Tables S1-2 show the raw data. Gene Ontology (GO) analyses of the identified Imp interactors revealed enrichment of proteins involved in several RNA metabolic pathways, including RNA localization, translation, gene expression regulation by silencing, and splicing (Fig. 1C, Fig. S4A). Proteins involved in oogenesis, regionalization, and cell maturation, processes that also depend on RNA localization, translational control, and tight regulation of gene expression, were also enriched among the GO terms associated with Imp interactors (Fig. 1C, Fig. S4A). The top 100 interactors include many RNA-binding proteins, and their association with Imp was affected by RNase treatment, suggesting that they may be part of BicD complexes through shared RNA targets (Fig. S3A-C). GO analysis also revealed that interactors whose presence in the Imp complex was reduced by RNase treatment were primarily enriched in RNA metabolic processes (Fig. S4B). Among the top 100 interactors, only the presence of Fmr1 and CG5726 was clearly not reduced in Imp complexes in the presence of RNase, suggesting a direct interaction with Imp (Fig. S3C). CG5726 is the *Drosophila* ortholog of human CTIF, a cap-binding complex-dependent translation initiation factor with no previously described function in *Drosophila*. Fmr1, which displayed slightly increased association with Imp in the presence of RNase, is an RNA-binding protein involved in RNA trafficking, translation, and neuronal excitability. In *Drosophila*, Fmr1 acts as a component of BicD/dynein transport complexes in neurons (Bianco et al., 2010), and its loss in humans causes the Fragile X syndrome.

Notably, several other BicD-interacting proteins, including Egl, Fmr1, PABP, Dhc, alongside the proteins involved in RNA localization in oogenesis, Stau and Sqd, were enriched in the Imp-GFP immunoprecipitations compared to the peptides detected in the Imp-GFP^Stop^ immunoprecipitations (Fig. 1D). These results confirm that Imp associates with BicD and BicD-associated proteins involved in RNA transport. We have previously shown that aside from its role in RNA transport, BicD also has a role in meiotic progression through interacting and properly localizing clathrin heavy chain (Chc) to meiotic spindles (Vazquez-Pianzola et al., 2022). Remarkably, Chc showed an enrichment in the presence of RNase in the Imp-GFP pull-downs (Fig. 1D). Furthermore, GO analysis revealed that in the absence of RNA, Imp is associated with factors involved in mitotic and meiotic cell cycle processes, chromosome segregation, spindle organization (Fig. S4C, D). These results further point to novel moonlighting functions of Imp in these processes.

### Imp is recruited to apically localizing RNAs in blastoderm embryos, but is redundant for RNA localization

Imp overexpression during oogenesis affects dorsoventral polarity and the localization of *grk* and *osk* maternal RNAs (Boylan et al., 2008; Geng and Macdonald, 2006). Furthermore, the Imp-binding sequences in *osk* mRNA are required for *osk* RNA anchoring, localization, and translational regulation (Munro et al., 2006). Nevertheless, *Imp* mutant germline clones do not block maternal mRNA localization during oogenesis, indicating a redundant role for Imp in mRNA localization in the ovarian germline (Boylan et al., 2008; Geng and Macdonald, 2006; Munro et al., 2006). Its involvement in the mRNA localization of BicD targets in early embryogenesis was not investigated and could be a potential cause for embryonic lethality. To test if Imp associates with mRNAs that are transported by BicD and Egl in the embryo, we injected a fluorescently labeled variant of the *hairy* (*h*) mRNA *(hSL1x3)*, which contains multiple copies of the Egl-BicD binding site and monitored Imp localization through the fluorescence of the Imp-GFP trap protein or antibodies against Imp. Once injected into the basal side of wild-type blastoderm embryos, these *h* transcripts are transported in a BicD/Egl/dynein-dependent manner to the cytoplasm apical to the nuclei (Bullock and Ish-Horowicz, 2001). Consistent with the presence of Imp in BicD complexes, we found that Imp co-localized with *hSL1x3* and BicD at the apical sites (Fig. 2Aa-b). Like BicD, Imp-GFP is seen apically of the nuclei (Fig. 2Ac) and upon basal injection of localizing RNAs, it hyperaccumulates with the injected RNAs apically of the nuclei (Fig. 2A.c-d, (Bullock and Ish-Horowicz, 2001)). Furthermore, inhibition of Imp activity by pre-injection of anti-Imp antibodies in wild-type embryos, as well as pre-injection of anti-GFP antibodies in Imp-GFP expressing embryos before injection of localizing *h* and *fushi tarazu* (*ftz*) RNAs, disrupted their apical transport (Fig. 2B-C). However, the localization of endogenous *ftz* and *h* RNAs remained unaffected in *Imp* mutant embryos generated by inducing mutant germline clones (GLCs) (Fig. 2D, Fig. S5A for information on *Imp* mutant alleles). Nevertheless, the inhibition of apical localization of injected mRNAs by preinjecting anti-Imp and anti-GFP antibodies suggests a mechanism where these antibodies sequester Imp-associated proteins necessary for the formation of a functional ribonucleoprotein transport complex. The finding that Imp interactors were enriched for RNA-binding proteins involved in RNA transport, stability, and translation control supports this notion (Fig. 1C-D, Fig. S3-4).

**Fig. 2.**
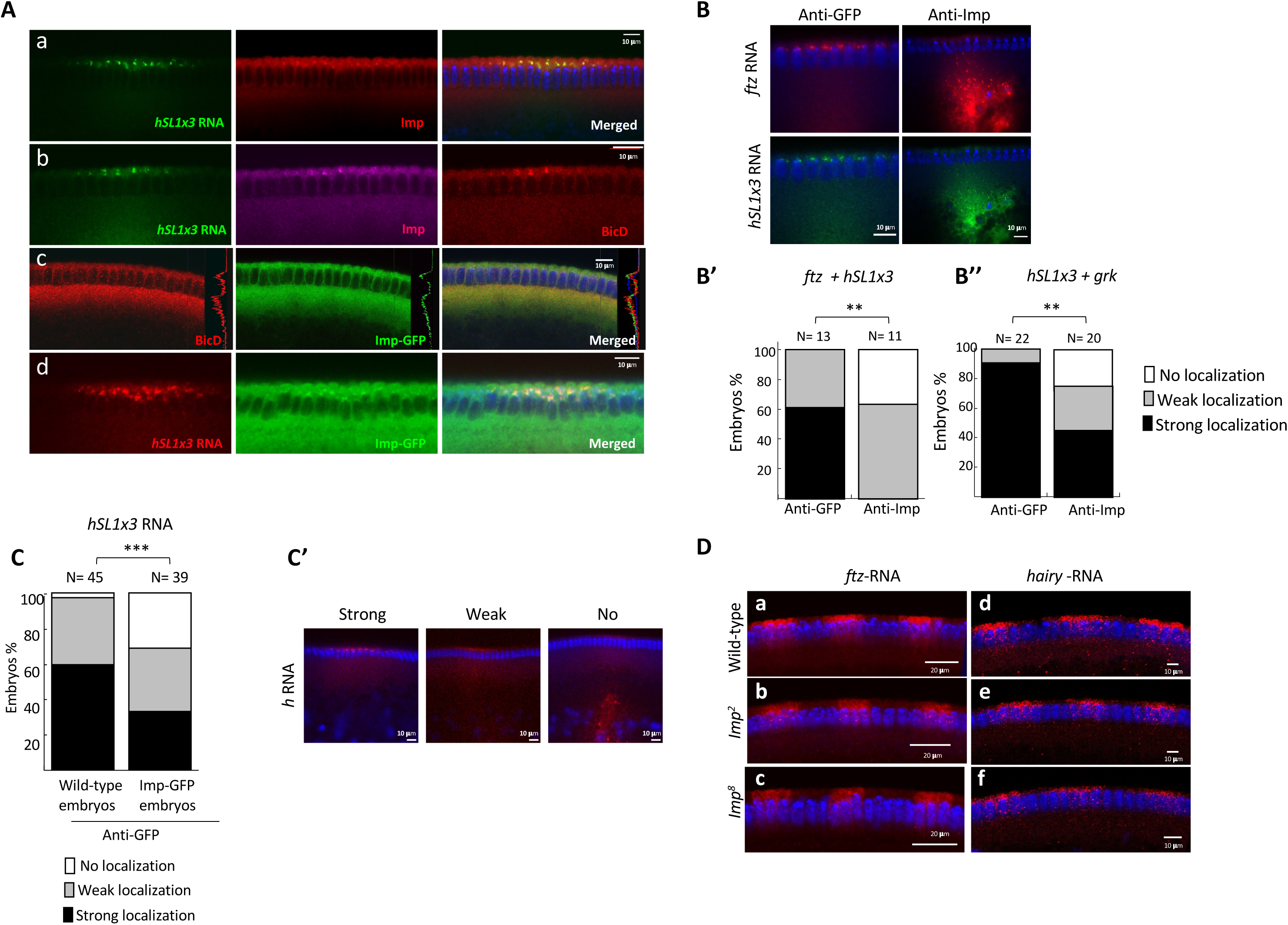
Co-recruitment of Imp and BicD to apically localizing *hairy (h)* RNA and Imp redundancy for endogenous mRNA apical localization. **(A,a)** Immunostaining to reveal the distribution of Imp (**a**, red) following injection of *hSL1x3* transcripts (green) into wild-type embryos. The merged image is on the right. **(A,b)** Imp (magenta) and BicD (red) protein distribution in wild-type embryos injected with localizing *hSL1x3* RNA (green). Imp and BicD are co-recruited to apically localizing *hSL1x3* RNA in this case bearing a poly-A tail (A_71_) that stabilizes the RNA. **(A,c)** Immunostaining of *Imp-GFP^G080^* embryos with anti-BicD antibodies (red). GFP signal is shown in green, and the merged image is seen on the right. Both proteins colocalize apically of the nuclei, although they also display a strong basal accumulation. **(A,d)** Distribution of Imp-GFP following injection with *h* transcripts into *Imp-GFP^G080^*embryos, Imp-GFP (green) is recruited to *hSL1x3* localized transcripts (red). The merged image is on the right. **(B-C).** Pre-injection of antibodies against Imp blocks apical localization of injected transcripts. **(B)** Distribution of injected (*ftz* (red) and *h* (green)) transcripts after pre-injection with anti-GFP (as control) or antibodies against Imp. Imp antibodies block apical colocalization and lead to the accumulation of injected RNA in the basal cytoplasm. **(B’)** Percentage of embryos displaying strong (black bars), weak (grey bars), or no localization (white bars) of injected *h* and *ftz* transcripts from the experiment shown in **B**. More embryos showed unlocalized RNAs upon injection with anti-Imp antibodies. Antibodies were injected 9-10 min before injection of a mixture of *h* and *ftz* RNAs. Embryos were fixed 6 min after RNA injection. (Distribution of staining intensity (Strong, Weak, None apical RNA localization) differed significantly between the two groups (Fisher’s exact test, 2×3 contingency table: χ²(2) = 9.57, p = 0.0083, **\*\***p < 0.01)**. B’’)** Similar experiment as in **B**, but a mixture of *hSL1x3* and *grk t*ranscripts was injected and embryos were fixed 9-12 min after RNA injection. Percentage of embryos displaying strong (black bars), weak (grey bars) or no localization (white bars) of the injected transcripts after injection of anti-GFP or anti-Imp antibodies into wild-type embryos. Upon pre-injection with anti-Imp antibodies, apical localization of injected transcripts was affected to a greater extent than upon injection with anti-GFP antibodies. Statistical significance was assessed using Fisher’s exact test on a 2×3 contingency table (χ²(2) = 9.74, p = 0.0077, **\*\***p < 0.01). **(C)** Percentage of embryos displaying strong (black bars), weak (grey bars), or no localization (white bars) of injected *h* transcripts after pre-injection of anti-GFP antibodies into wild-type or *Imp-GFP^G080^*expressing embryos. Antibody was injected 10-17 min before RNA injection, and embryos were fixed 6 min later. 30 % of *Imp-GFP^G080^* embryos showed unlocalized *h* RNA compared with only 2 % observed in wild-type embryos. Distribution of staining intensity (Strong, Weak, No) differed significantly between groups (Fisher’s exact test on a 2×3 contingency table: χ²(2) = 14.14, p = 0.00085, ***p < 0.001). **(C’)** Examples of *Imp-GFP^G080^* embryos pre-injected with anti-GFP antibodies before injection with *h* RNA (red). Examples scored as strong, weak, and no *h* localization are shown. (D) *In situ* hybridization to whole-mount wild-type (a, d), *Imp^2^* (b, e), and *Imp^8^* (c, f) embryos using digoxigenin-labeled *ftz* (a-c) or *h* (d-f) anti-sense RNA probes (red signals). The embryos are also stained with Hoechst (blue) to visualize the DNA. No defects in apical RNA localization of pair-rule transcripts were observed in embryos devoid of maternal *Imp.* The same results were obtained with the *Imp^7^* allele (not shown).

These observations collectively suggest that while Imp–RNP complexes are crucial for proper RNA transport, Imp itself is not an essential component, or adaptation to the loss of Imp begins early in development, indicating functional redundancy of Imp during early stages.

### Transcriptomes of *Imp* mutant embryos display dysregulation in stem cell versus differentiation pathways

Embryos devoid of maternal *Imp* during early development showed no defects in pair-rule mRNA localization (Fig. 2D). However, cuticle preparations of embryos devoid of maternal Imp (derived from *Imp* germline clones (GLCs)) revealed predominant defects in anterior development, consistent with previous reports (Boylan et al., 2008)(Fig. S5B). These defects resulted in defective dorsal closure and seem to happen during germband retraction or head involution (Fig. S5C).

To deepen our understanding of the role of maternal *Imp* throughout embryogenesis, we conducted a detailed RNA transcriptome analysis comparing *Imp^+^*embryos (*y w*) with embryos derived from induced germline clones of two distinct *Imp* null mutant alleles, namely *Imp^2^* and *Imp^8^* (refer to Fig. S5A and the methods for allele descriptions). Our study involved total RNA sequencing of embryos aged 0-4 hours (before visible phenotypes emerged) and 0-8 hours (when defects in germband retraction started to appear) for *Imp* mutants and control *Imp^+^*embryos. RNA sequencing was performed on biological triplicates. Both *Imp* mutants exhibited similar disparities in gene expression compared to *Imp^+^*embryos, causing them to cluster together in the heat map of gene plots (Fig. 3, raw differential expression (DE) counts data in Tables S3-S6). Among the top misregulated genes in the absence of *Imp,* we observed a significant prevalence of upregulated genes over downregulated ones, suggesting that Imp primarily functions as a repressor of gene expression (Fig. 3). Additionally, GO analysis of the most differentially expressed genes in 0-4-hour embryos revealed downregulation of genes associated with translation, RNA metabolism, RNA splicing, and cell growth. Whole embryo immunostaining and cuticle preparations revealed that *Imp* embryos are smaller compared to wild-type embryos (Fig. S5B-C), reinforcing the role of *Imp* in embryo growth. Interestingly, this size reduction is reminiscent of the ∼40% smaller size observed in *Imp1*-deficient mice, and it indicates a proportional decrease in most organ sizes (Hansen et al., 2004). In contrast, genes linked to morphogenesis, tube morphogenesis, muscle development, differentiation, and negative regulation of cellular biosynthesis and transcription were upregulated (Fig. 4A). Similarly, 0-8-hour embryos displayed upregulation of genes linked to tube morphogenesis and neuronal projection development and abnormally branched trachea, indicative of defects in morphogenesis, particularly during tube formation and branching (Fig. 4C, C’). The roles of *Imp* in activating genes involved in cell growth and repressing genes involved in differentiation suggest that premature differentiation causes the defects in gastrulation and differentiation. *Imp* may thus play a role in coordinating cell numbers with differentiation—potentially by repressing differentiation until normal cell numbers are achieved.

**Fig. 3.**
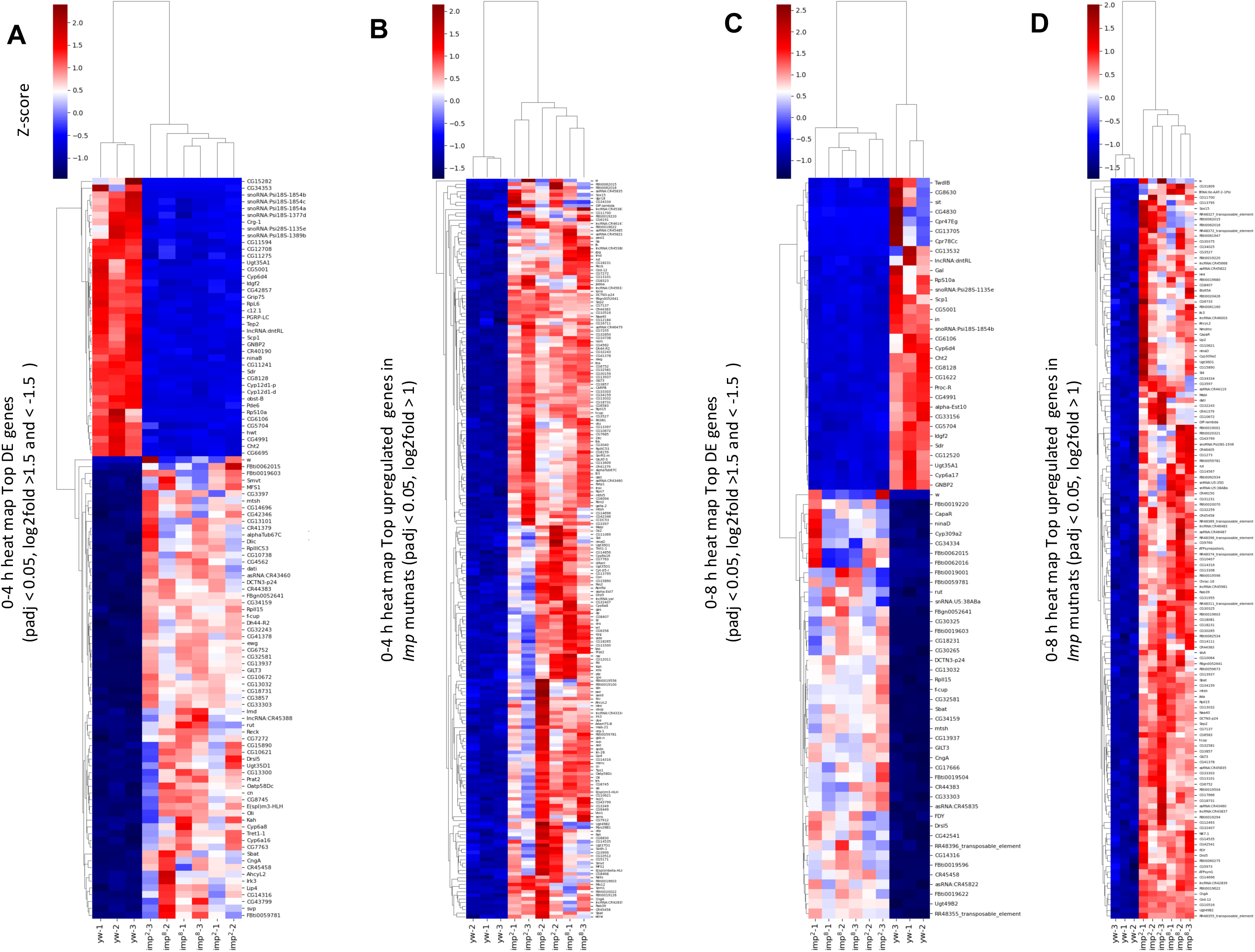
Transcriptome profile of embryos lacking maternal *Imp.* Total RNA sequencing was performed in triplicates on embryos aged 0-4 hours (A-B) and 0-8 hours (C-D) derived from the induction of *Imp^2^* and *Imp^8^*germline clones (GLCs), alongside control wild-type embryos (yw). Heat maps display the top differentially expressed (DE) genes (A, C) or only the top upregulated genes (B, D) in the mutants compared to wild-type embryos. Selected cutoffs are indicated. Both *Imp* mutant embryos cluster closely together in the heat maps.

**Fig. 4.**
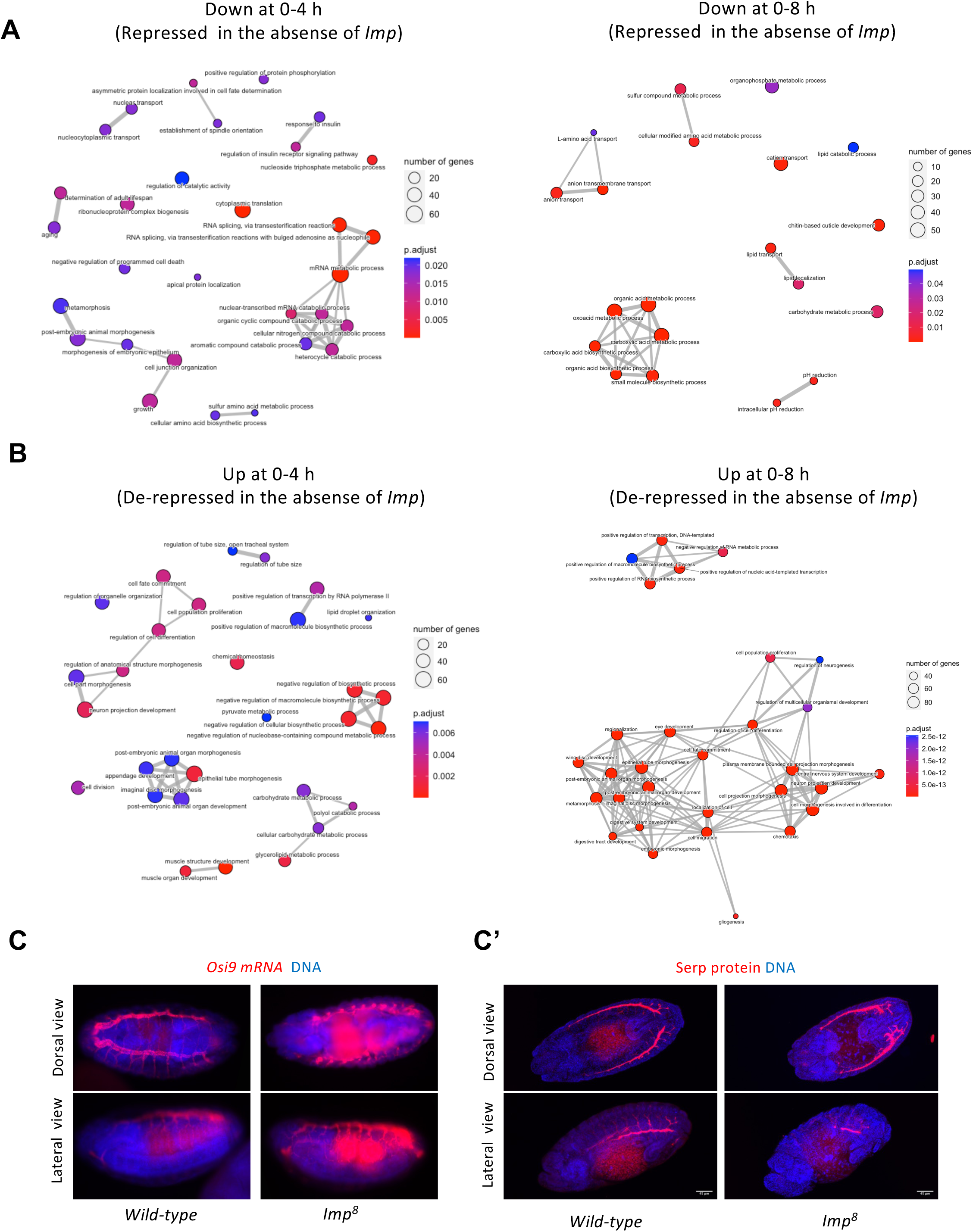
Gene Ontology Analysis (GO) of deregulated RNAs in *Imp* mutants reveals functional links to observed phenotypes. **(A-B)** GO analysis was performed on the list of genes found to be significantly (Padj < 0.02) differentially expressed in both *Imp^8^* and *Imp^2^* mutants in the 0-4 h and 0-8 h old collections, compared to controls. The GO analysis was conducted separately for downregulated (A) and upregulated (B) genes. Cutoff for upregulated genes: log2FoldChange > 0. Cutoff for downregulated genes: log2FoldChange < 0 in 0-4 h old embryos and < -0.04 in 0-8 h old embryos. **(C)** mRNA *in situ* hybridization against *Osiris9 (Osi9)* mRNA (red), a gene expressed in the tracheal system. **(C’)** Embryos immunostained for the chitin deacetylase Serpentine (Serp) protein, a marker of the tracheal lumen (Hansen et al., 2004). *Imp* mutant embryos display abnormally branched trachea and a smaller body size. Embryos were counterstained with Hoechst to visualize the nuclei.

### Imp associates with mRNAs involved in pathways governing cell growth, cell cycle, stem cell regulation, and differentiation

We then conducted RIP-seq analyses to pinpoint RNAs interacting with Imp particles. Immunopurification of Imp-RNA complexes was performed using *Imp-GFP^G080^* embryo extracts and anti-GFP antibodies. As a mock control, we also sequenced RNAs immunoprecipitated with anti-GFP antibodies from extracts of embryos expressing the non-functional N-terminal version of Imp fused to GFP (*Imp-GFP^Stop^*) (Fig. 5; raw data in Table S7; Fig. S2). Additionally, total RNA sequencing was performed on the input extract utilized for the Imp::GFP pulldown as an additional control (Fig. 5A-B; raw data in Table S8). A high overlap was observed for transcripts enriched in the Imp-GFP^G080^ IPs compared to the input RNA control and the same IP compared to the compared to the *Imp-GFP^Stop^* mock IP control (Venn diagram, Fig. 5A). The heat map illustrating the top RNAs enriched in the Imp::GFP^G080^ IP vs Imp::GFP^G080^ input RNA and the *Imp-GFP^Stop^* mock IP control show that many of these RNAs encode ribosomal proteins, long non-coding RNAs (lncRNAs) and TE RNAs (Fig. 5B). Further analysis of Imp-associated RNAs (from the genes in the overlap in Fig. 5A) through GO enrichment highlighted terms closely aligned with those RNAs differentially expressed in the *Imp* mutants (Fig. 5C–Fig.4A-B). This included genes linked to translation, RNA splicing, and genes involved in regulation of tube size, the tracheal system and embryonic development.

**Fig. 5.**
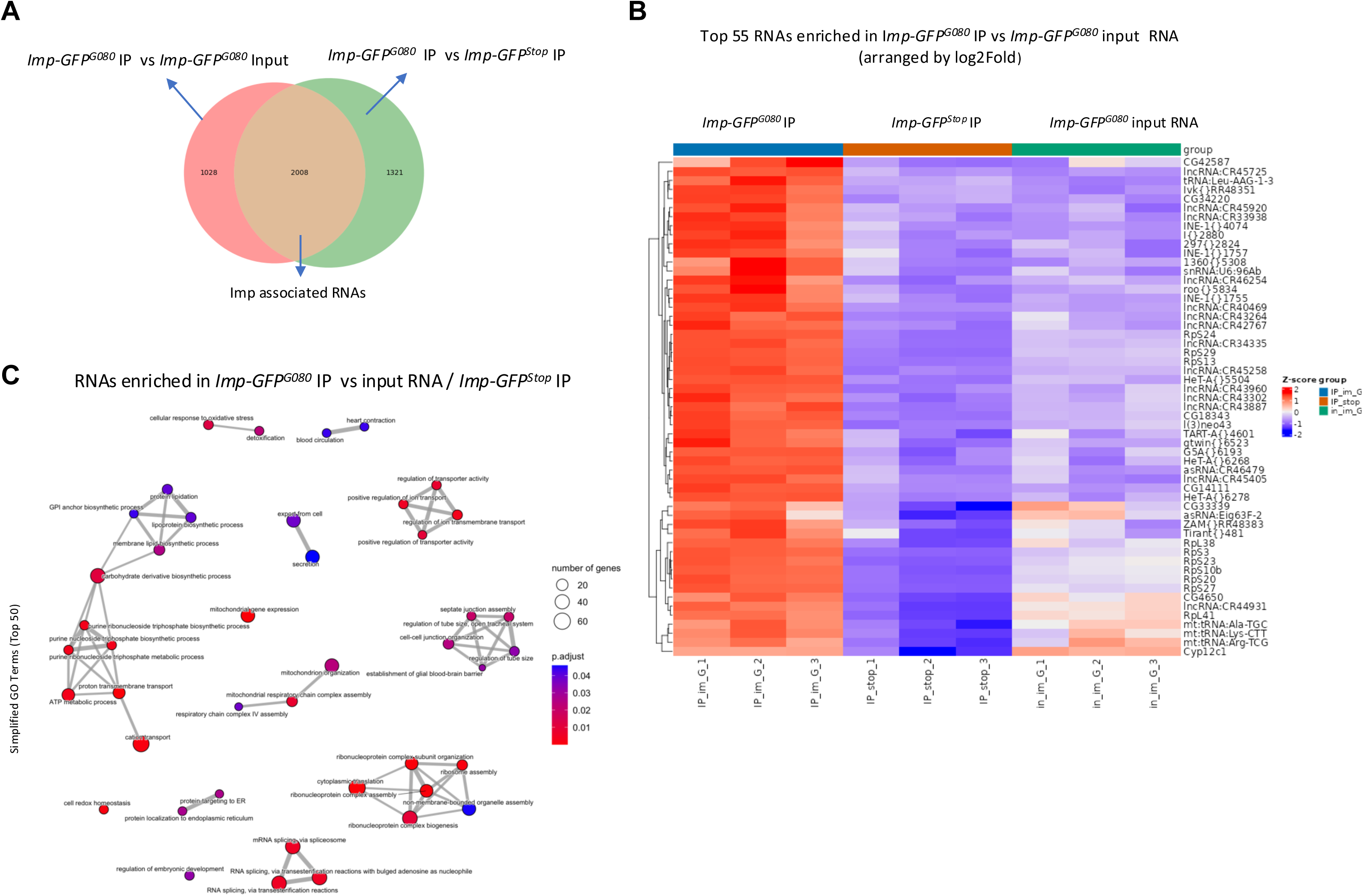
Identification of RNAs present in Imp-RNP complexes. **(A)** Venn diagram showing the overlap between the genes found enriched in *Imp*-GFP IPs compared to the mock control IPs and those found enriched in the *Imp*-GFP IPs compared to the input RNAs. A cutoff of log2FoldChange > 0 and Padj < 0.05 was used for this graph. A high degree of overlap was observed. **(B)** Heat map illustrating the top 55 differentially expressed (DE) genes with the highest absolute log2FoldChange difference between *Imp*-GFP and the mock IPs (Padj < 0.05). The heat map reveals similar expression patterns between the input and mock control IPs (*Imp-GFP^Stop^*) compared to the *Imp*::GFP IPs. **(C)** GO enrichment analysis of *Imp*-associated RNAs. The RNAs selected as enriched using the two different controls depicted in the Venn diagram in **(A)** are shown.

### Imp associates with TE mRNAs and represses them

Unexpectedly, in both 0-8- and 0-4-hour-old *Imp* mutant embryos, a substantial increase in RNAs originating from transposable elements (TE) was evident (Fig. 3B-D, raw data in Tables S3-S6). TE RNAs were prominently overrepresented in the *Imp* mutants compared to wild-type controls, and the increase was even more pronounced in the 0-8-hour embryo collections, indicating that TEs were derepressed in the mutant embryos (Fig. 6A). Our standard RNA sequencing pipeline captured this difference in expression, likely due to polymorphisms across various transposon insertions. Subsequent utilization of a specialized RNA sequencing pipeline, targeting transposable element expression, further validated and strengthened the observed transposon de-repression in *Imp* mutant embryos (Fig. 6B, B’) (Pogorelcnik et al., 2018). Several transposons were highly upregulated in the *Imp* mutant embryos. Furthermore, the top list of RNAs that co-immunoprecipitated with Imp-GFP^G080^ was also notably enriched with TE RNAs and non-coding RNAs (Fig. 6C). These enrichments are real since the representation of these biotypes was increased only in the list of RNAs enriched in the Imp-GFP^G080^ IP and not in the RNAs not found enriched in the IPs (Fig. 6C). TE RNA sequencing mapping also identified specific TE enriched in the Imp-GFP ^G080^ IPs (Fig. 6D). This correlates with biotypes overrepresented in the RNAs found to be upregulated in *Imp* mutant embryos in the transcriptome analysis (Fig. 6A). Accordingly, a proteomic analysis of 0-4-hour-old embryos unveiled the presence of peptides specific to TE-encoded proteins solely in *Imp* mutant embryos. Peptides corresponding to *roo* elements (|Q8I7Q1), Envelop (Env, Q5K600) protein from the *Tirant* transposon, and the Gag protein (O01350) from the *Burdock* transposable element were specifically detected in *Imp^8^* and *Imp^2^* embryos but were absent in their *Imp^+^*counterparts (Table 1).

**Fig. 6.**
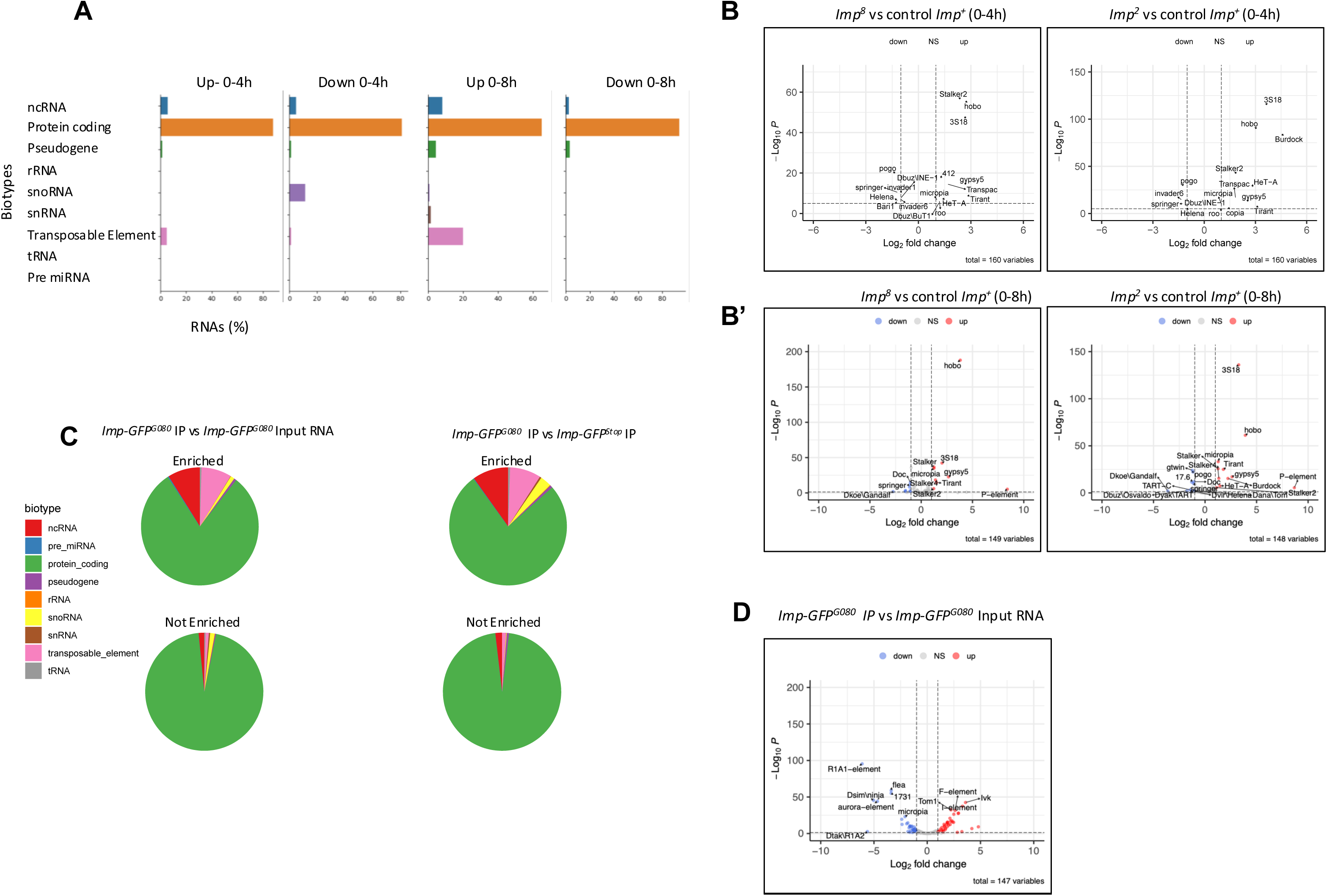
Transposable element (TE) mRNAs and ncRNAs are upregulated in *Imp* mutant embryos, and they also associate with Imp. **(A)** Percentage of RNA biotypes among the genes found to be either upregulated or downregulated in both *Imp* mutants compared to control embryos. **(B)** RNAs found to be differentially expressed between *Imp^2^* and *Imp^8^* mutants on one side, and control embryos in the 0-4 h collections were specifically mapped to TEs. Volcano plots are shown. **(B’)** Same as **(B)**, but in 0-8 h old embryos. **(C)** Percentage of RNA biotypes observed among genes found enriched in the *Imp*-GFP IP compared to the two different controls described in Fig. 5 (input and mock IP). The percentage of biotypes among RNAs not enriched in the IPs is also shown as a control. ncRNAs and TE RNAs are the most enriched RNAs in Imp-GFP pull-downs. **(D)** Volcano plot showing the RNAs pulled down with Imp-GFP specifically mapped to TEs.

**Table 1.**
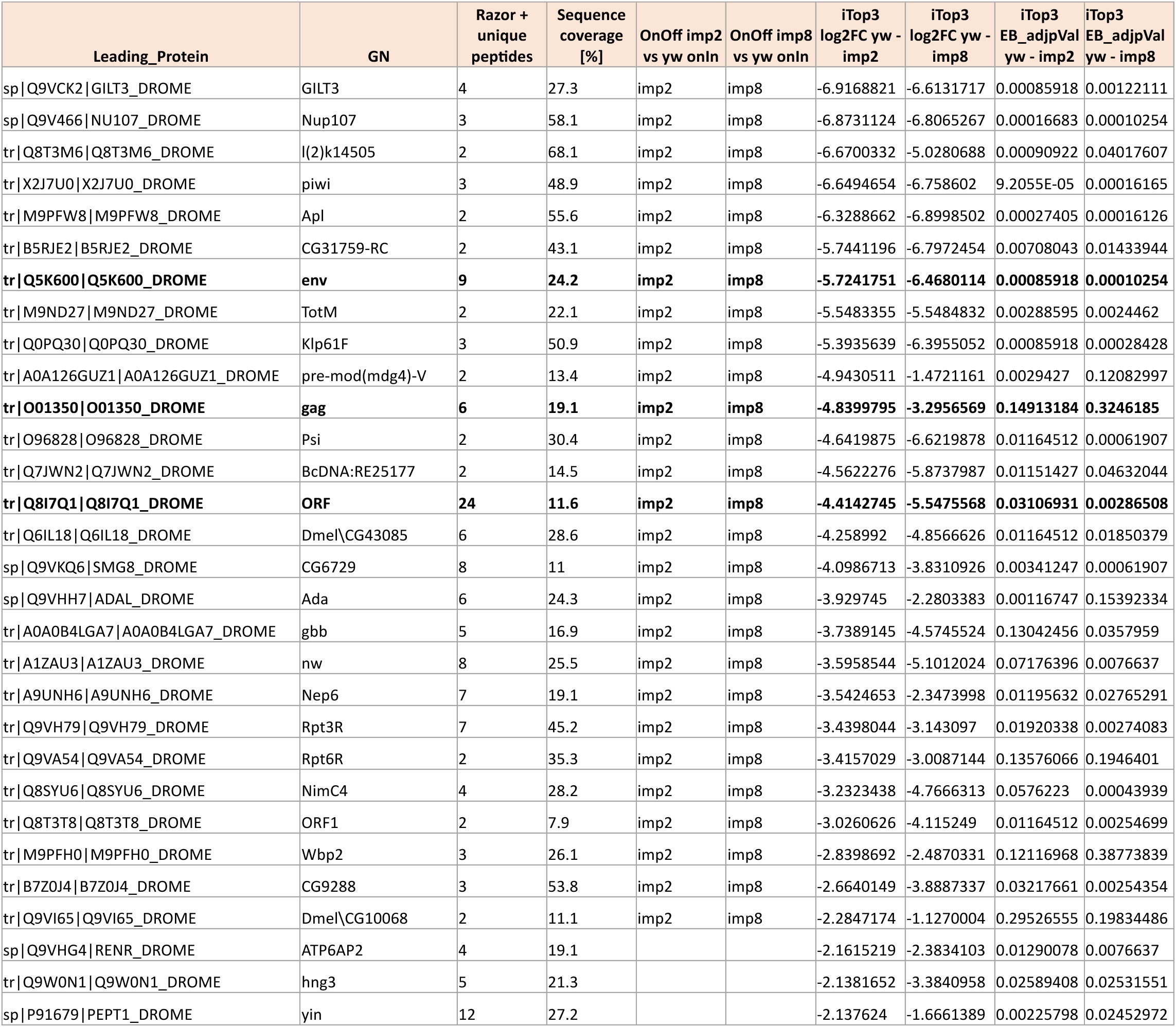
Transposable Element–Encoded Proteins Are Derepressed in *Imp* Mutants. Top 30 up-regulated proteins in *Imp* mutant embryos. Embryos collected 0–4 hours after egg laying from wild-type (*yw*) and *Imp*-depleted backgrounds—generated using germline clones (GLCs) of the *Imp^2^* and *Imp8* alleles—were analyzed by mass spectrometry. The 30 proteins with the highest log₂ fold change in both *Imp* mutants compared to wild-type samples are shown. Proteins were selected based on a p-value cutoff of 0.05 or if they were exclusively detected in one condition but not the other (as indicated in the “OnOff” column, which specifies the condition in which the protein was detected). TE encoded proteins are highlighted in bold.

Staining *Imp* mutant embryos with an antibody against the Gag/pol protein encoded by the Copia transposable element detected distinct signals in specific cells or groups of cells (Fig. 7A). Conversely, no or only minimal positive signals were observed in wild-type embryos. Copia^gag^ expression emerged as early as stage 12 of embryogenesis in *Imp* mutants, becoming notably prevalent in yolk cells by stage 14. Western blot analysis confirmed Copia overexpression in *Imp* mutant embryos (Fig. 7B).

**Fig. 7.**
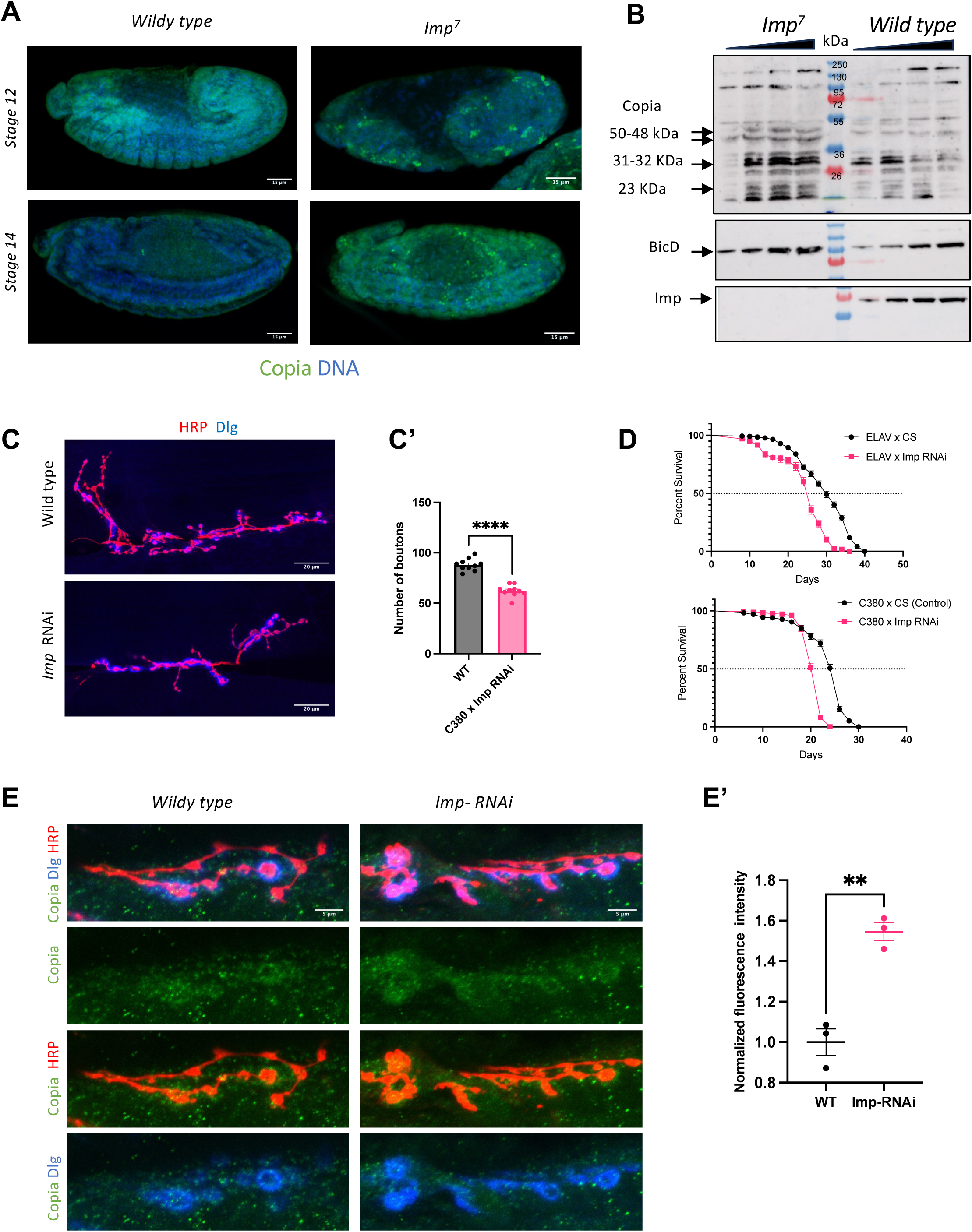
Copia protein is upregulated in *Imp* mutant embryos and at NMJs with reduced *Imp* levels. Embryos lacking maternal *Imp* (*Imp^7^* GLCs) and wild-type embryos were subjected to immunofluorescence staining using anti-Copia antibodies (green signal). DNA stained with Hoechst is shown in blue. Stage 12 and stage 14 embryos are shown. In wild-type embryos, the Copia signal remains at background levels, similar to staining with only the secondary antibodies. In contrast, *Imp* mutant embryos show clear and specific expression of Copia in individual cells and cell clusters. **(B)** Copia protein was detected by Western blot, too. Anti-BicD was used as a loading control. Imp protein is absent in *Imp^7^* mutants. Copia is expressed in different bands (indicated by arrows), which may represent various proteolytically cleaved fragments with a main band around 50 KDa corresponding to the Gag protein. **(C)** Control and *Imp* RNAi expressing larval NMJs stained for Horse Radish Peroxidase (HRP) and Disc Large (DLG) to trace the motor neurons and the postsynaptic machinery, respectively. Knockdown of *Imp* causes a reduction in bouton numbers. **(C’)** Quantification of bouton numbers. **(D)** Knockdown of *Imp* using either the pan-neuronal driver *elav*-GAL4 or the motor neuron-specific driver *C380*-GAL4 causes a reduction in lifespan. **(E)** Representative confocal images of NMJs stained to detect Copia Gag (green), HRP (red, motoneuron marker), and Dlg (blue, postsynaptic marker) in wild-type larvae (C380-Gal4/CantonS) and upon knocking down (KD) *Imp* in the presynaptic compartment with C380-Gal4 RNAi. **(E’)** *Imp* KD causes an increase of α-Copia Gag signal at larval NMJs compared to wild-type NMJs.

The observed misregulation of genes involved in the development of neuronal projections, led us to examine the larval neuromuscular junction (NMJ). Knocking down *Imp* at the larval NMJ by expressing *Imp* RNAi under the control of the motor neuron-specific driver C380 reduced the number of synaptic boutons (Fig. 7C, C’). Although these flies survive to adulthood, they show a reduced lifespan, confirming a critical role for *Imp* in neurogenesis and its importance for fly survival (Fig. 7D). We found that this phenotype correlated with increased expression of Copia, which appeared condensed in and around the larval NMJs following *Imp* RNAi knockdown (Fig. 7E-E’). Altogether, this points to a role for Imp in repressing Copia expression at larval synapses and during embryogenesis.

### Loss of *Imp* neither impairs nuclear export of piRNA precursors nor piRNA production

Because TEs were de-repressed in *Imp* mutants, we investigated whether *Imp* affects the piRNA pathway, the primary defense against transposons. The regulation of transposon families expressed during early development depends on maternally deposited piRNAs and Piwi proteins (Fabry et al., 2021). piRNA precursors, originating from the *Flam* and *42AB* loci, are pivotal in suppressing transposable elements in both the soma and germline of *Drosophila*, particularly targeting retrotransposons like the *gypsy* family, which exhibited heightened expression in *Imp* mutants. Although Imp’s orthologs have suggested roles in nuclear RNA export (Oleynikov and Singer, 2003; Pan et al., 2007), we found no evidence indicating that *Imp* is needed for the production or nuclear export of these piRNA precursors (Fig. S6A-B). We further investigated whether *piwi* RNA production was compromised in *Imp* mutant embryos by sequencing the small RNAs in control and *Imp* mutant embryos (Fig. S6C-D). The percentage of total small RNAs mapping to TEs was not affected in embryos devoid of maternal *Imp* (Fig. S6C). Although the levels of piRNAs mapping specifically to some de-repressed TEs were slightly reduced in the *Imp* mutants (Tirant, HET-A, Stalker-2, microcopia), this expected inverse correlation did not generalize to the other TEs de-repressed in *Imp* mutants (such as Copia) (Fig. S6D). We next investigated whether the ping-pong amplification cycle was affected by *Imp* loss by quantifying the fraction of piRNA read pairs with 10-nt 5′ overlaps between sense and antisense reads. Across all samples, we found only a small subset of transposon-mapping piRNAs with such 5’ overlaps (Fig. S6E). For these piRNAs, the percentage of sense and antisense pairs with a 10-nt overlap did not differ between wild-type and *Imp* mutant samples. These observations suggest that *Imp* does not influence the ping-pong amplification pathway and piRNA production. However, since the number of piwi RNA reads detected in embryos was very low, they may have masked the role of *Imp* in *piwi* RNA production if it functions earlier during oogenesis.

### Imp associates with proteins involved in gene silencing

Finding no effects on piRNAs, we closely examined the proteins present in complex with Imp to learn more about the mechanism of TE regulation by *Imp* in embryos (Fig. S3, Fig. 1C). GO analyses of Imp interactors revealed a strong association with proteins involved in negative regulation of gene expression and gene silencing (Fig. 1C, S3A). Notably, in the Imp-GFP^G080^ immunoprecipitates, several components of the RNA processing machinery involved in mRNA and transposon silencing were enriched (Fig. 8A). Key players in the piRNA production and amplification pathway—such as Piwi, Spn-E, Tdrd3, Hop, Aub, Ago3, and Cbp80—were identified (Fig. 8A).

**Fig. 8.**
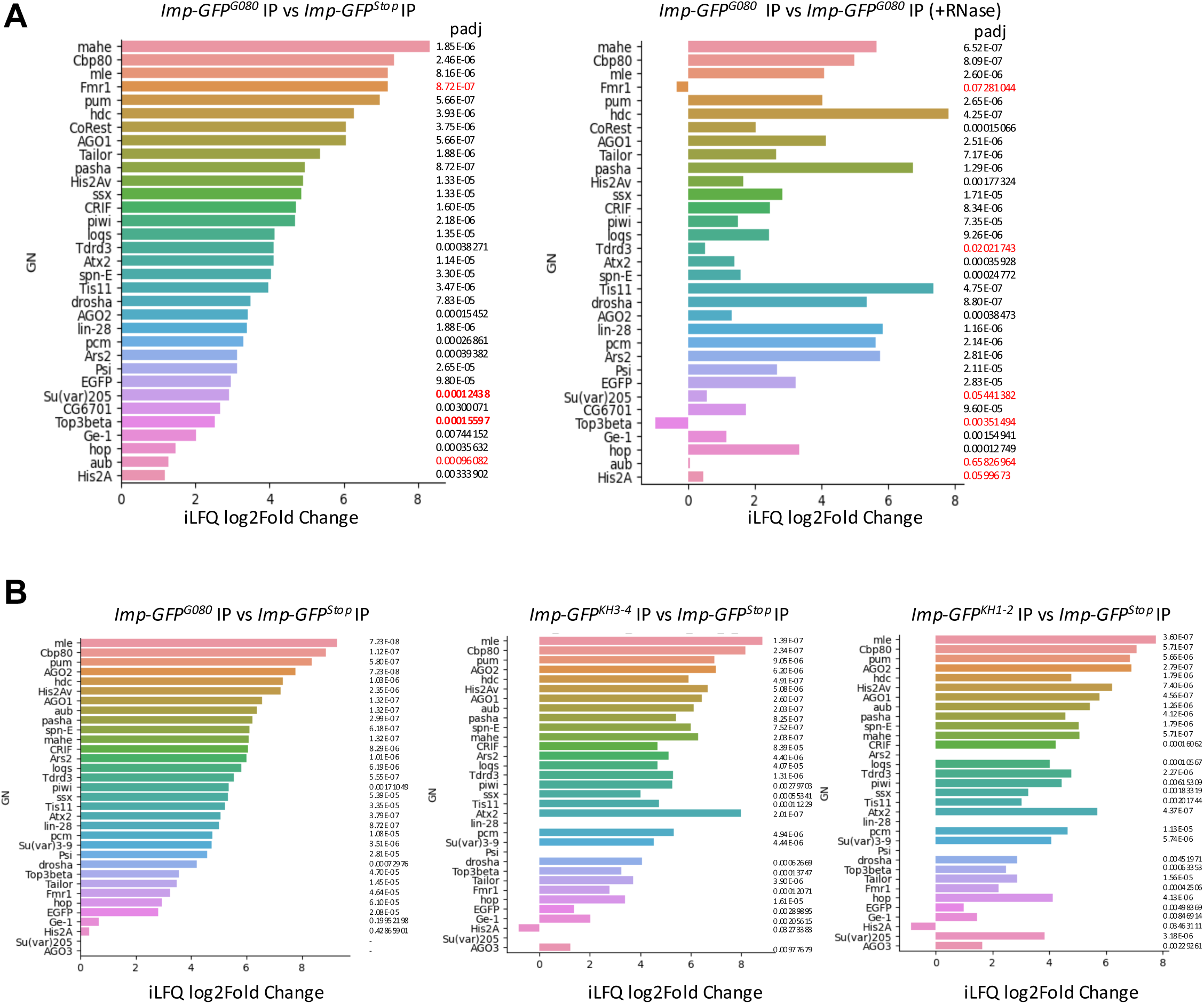
*Imp* associates with proteins involved in TE silencing. The log2FoldChange enrichment of selected proteins, known to be involved in RNA silencing pathways, is shown for different versions of Imp-GFP proteins compared to the mock controls. The corresponding padj values of the enrichment are also depicted. **(A)** Enrichment between the peptides found in the *Imp-GFP^G080^* IPs over the ones found in the control IPs (*Imp-GFP^Stop^ IP*) (left panel) or the *Imp-GFP^G080^* IPs done in the absence and presence of RNase are shown (right panel). **(B)** Enrichments between the peptides found in the *Imp-GFP^G080^* IPs and the KH mutant versions of Imp-GFP (Imp-GFPKH3-4 and Imp-GFPKH1-2), compared to the mock IPs (*Imp-GFP^Stop^ IP*), are shown. Data from A and B represent different experiments, and pull-downs were performed in biological triplicates.

Other factors involved in heterochromatin formation and transposon silencing remained associated with Imp even in the presence of RNase during the immunoprecipitation experiments, namely Fmr1, Top3beta, Tdrd3, Su(var)205/HP1, Aub, and His2A, indicating a direct binding of Imp with some of them, regardless of shared RNA or TE mRNA targets (Fig. 8A). However, the pattern of histone H3 lysine 9 trimethylation (H3K9me3) and Hp1 localization in *Imp* mutant embryos did not show any clear visible disruption (Fig. S7A).

### Piwi protein is not properly localized to the nuclei of *Imp* mutant nurse cells

Our proteomic analysis of *Imp* mutants revealed that Piwi, a key regulator protein of TE expression, was upregulated (Table 1). In *Imp* mutants, Piwi appears to be present during embryogenesis and enriched in all blastoderm nuclei from early embryogenesis on (Fig. S7B). Aside from its enrichment inside all blastoderm nuclei, Piwi appeared concentrated in dot-like structures in the heterochromatin. This pattern was not as strong in some *Imp* embryos derived from *Imp* GLCs, but the differences were subtle and below our threshold of detection (Fig. S7B). However, examination of the maternal germline revealed a striking mislocalization of Piwi protein in *Imp* mutant ovaries (Fig. 9A). In wild-type ovaries, Piwi protein localizes to the nuclei of both nurse cells and follicle cells, showing higher expression in the nurse cells compared to a lower signal observed in the follicle cells. In contrast, in most *Imp* mutant egg chambers, Piwi accumulates at lower levels in the nurse cells than in the follicle cells or fails to accumulate at all in the nurse cell nuclei (see Fig. Legend for quantification), while Piwi still showed nuclear localization in the surrounding follicle cells, which retained normal Imp expression (Fig. 9A). The strong phenotype in the germline, compared to the subtle one in embryos, may be because zygotic expression of other factors in the embryo partially compensates for or masks the maternal defect. Localization of Ago1, a regulator of the miRNA pathway, was not affected in *Imp* mutants, consistent with Imp having a specific role in the piRNA pathway independent of the miRNA pathway (Fig. 9B).

**Fig. 9.**
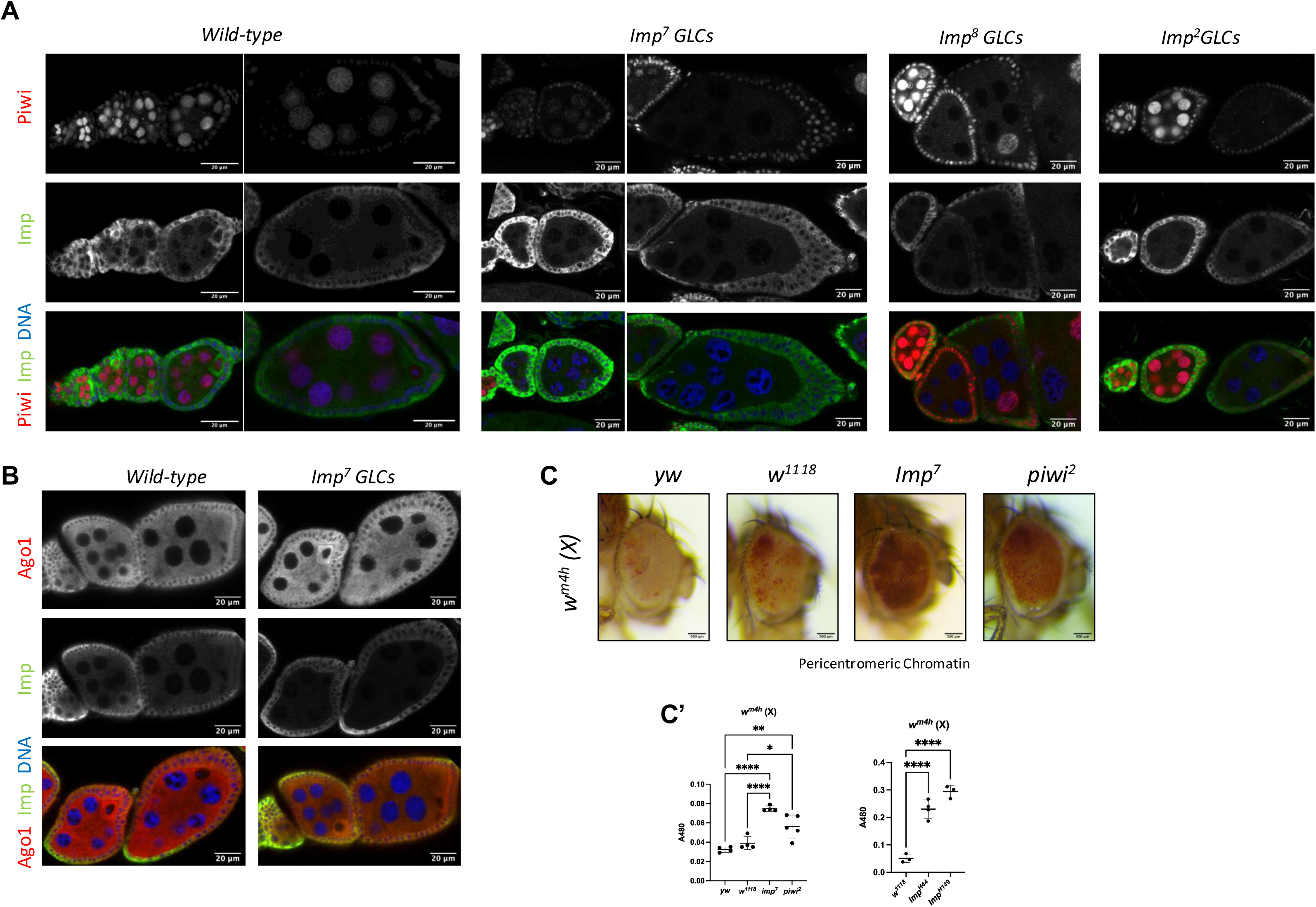
*Imp* is required to maintain Piwi localization in the germline during oogenesis and for transcriptional silencing by position-effect variegation (PEV) in the soma. **(A)** Piwi presence in stage 6 and 8 egg chambers was monitored by acquiring z-stack images. Piwi was localized to nurse cell nuclei in most wild-type egg chambers in two independent analyses (96.8%, n=30; 100%, n=25). In *Imp-*mutant GLCs, Piwi failed to localize in most nurse cell nuclei. Specifically, 84% of *Imp^8^* GLCs (n=25), 70.8% of *Imp^2^*, and 67.7% of *Imp^7^* GLC egg chambers (n=31) that did not express *Imp* in the germline, showed nurse cell nuclei where Piwi was not enriched, indicating that *Imp* is required to maintain nuclear Piwi during these stages in the germline. In these *Imp* germline mutant egg chambers Piwi still shows normal localization to the Imp-expressing follicle cell nuclei. **(B)** Ovarioles devoid of *Imp* in the germline and wild-type ovarioles were stained for Ago1 (red) and Imp (green). Ago1 localization seems not affected in *Imp* mutant egg chambers. **(C)** Representative images of eye pigmentation observed in the indicated genetic backgrounds. The *wm4h* white line reporter in the pericentromeric X chromosome region was used. *Imp^7^*and the *Imp^H44^* and *Imp^H149^* excision lines generated by mobilization of a different P-element were all defective in transcriptional silencing by PEV, similar to *piwi* mutants. **(C’)** Eye pigment quantification for the indicated genotypes was measured by absorbance at A480 nm.

### *Imp* mutants are defective in transcriptional silencing by position-effect variegation (PEV)

In *Drosophila,* components of the RNA-induced silencing complex (RISC) and the piRNA pathway, including Piwi, act as heterochromatin regulators and are implicated in Position Effect Variegation (PEV) (Lee et al., 2018). Therefore, we investigated whether Imp functions similarly using a standard PEV assay in the eye, a somatic tissue (Fig. 9C). In this assay, a reporter encoding the *white* gene inserted near heterochromatin (*wm4h*), results in variegated red eyes due to heterochromatin spreading and subsequent transcriptional silencing in some cells but not others. Like *piwi* mutants, females carrying the *wm4h* reporter but displaying reduced Imp levels due to heterozygosity for *Imp* alleles (*Imp7/+*, *ImpH44/+*, or *ImpH149/+*) showed an increased *mini-white* gene expression in their eyes compared to females with normal Imp levels (*Imp ±/+* background) (Fig. 9C-C’, Fig. S8). This suggests that like Piwi, Imp regulates transcriptional silencing by position-effect variegation (PEV).

## DISCUSSION

This study provides insights into the molecular mechanisms by which maternal Imp (IGF2BP) influences early embryonic development in *D. melanogaster*. Females with mutant *Imp* germline clones display normal ovaries, but their embryos do not develop, highlighting the critical role of maternal *Imp* for embryogenesis. Previous research suggested a role for Imp in mRNA localization during oogenesis, though this role appeared redundant (Boylan et al., 2008; Munro et al., 2006). Imp is also involved in neuronal RNP assembly and granule dynamics, regulating axonal remodeling (Vijayakumar et al., 2019). Immunoprecipitation, mass spectrometry, and yeast two-hybrid analyses identified Imp as part of BicD complexes, linking Imp to the BicD/dynein transport machinery in embryo extracts. Notably, BicD interacted with the Imp carboxy terminus which contains the Prion Like Domain (PLD), identified as critical for promoting the motility of axonal Imp granules (Vijayakumar et al., 2019). This domain is also essential for efficient Imp localization to axons and appears to be responsible for recruiting molecular motors. Despite Imp’s recruitment to apically localizing RNAs in blastoderm embryos, endogenous mRNA localization remained unaffected in *Imp* mutants. These observations support the notion that germline *Imp* is not essential for the apical localization of these mRNAs, and it might be redundant during *Drosophila* oogenesis and early embryogenesis for RNA transport, a conclusion that was also reached by others (Geng and Macdonald, 2006b; Munro et al., 2006) However, inhibition of Imp by pre-injection of anti-Imp antibodies disrupted the apical transport of injected mRNAs, suggesting that antibodies against Imp may prevent the localization of RNAs by blocking the formation of functional RNP localizing particles containing Imp and Imp-binding proteins. Indeed, Gene Ontology (GO) analysis of the identified Imp interactors showed an enrichment of proteins involved in RNA localization. Imp appears to interact directly with Fmr1, a protein known to associate with BicD (Bianco et al., 2010) and be involved in RNA transport and translational control (Prashad and Gopal, 2021). GO analysis revealed that many Imp interactors, which do not depend on RNA for their interaction, are factors involved in mitotic cell cycle processes, chromosome segregation, spindle organization, and nucleocytoplasmic transport. For example, Chc, a known BicD interactor involved in cell cycle regulation, was associated with Imp when RNA was depleted in the extracts. Remarkably, although the role of the Imp paralogs in nucleocytoplasmic transport has been demonstrated (Hüttelmaier et al., 2005; Oleynikov and Singer, 2003; Pan et al., 2007) their potential moonlighting role in cell cycle regulation requires further investigation.

Transcriptome analysis of *Imp* mutant embryos revealed significant dysregulation of genes associated with cell growth, differentiation, and RNA metabolism, along with upregulated TE RNAs. Genes involved in translation, RNA metabolism, RNA splicing, and cell growth were downregulated in *Imp* mutant embryos, suggesting that *Imp* positively regulates the expression or stability of these transcripts. In contrast, genes associated with morphogenesis, tube development, muscle development, neurogenesis, and differentiation were upregulated, indicating that *Imp* normally represses their expression at this stage of embryogenesis. The RNAs present in the purified Imp-RNP complexes displayed similar GO terms. Several of the phenotypes observed in *Imp* mutant embryos—such as smaller size compared to their *Imp*⁺ counterparts, defects in tracheal branching, incomplete dorsal closure in embryos, and defective larval NMJ formation—correlate with the role of *Imp* in controlling the expression of genes involved in molecular pathways that regulate stem cell function and differentiation.

TE-encoding mRNAs were also detected in Imp-RNP complexes, and both proteomic analysis and immunostaining confirmed the upregulation of TE-encoded proteins in *Imp* mutant embryos. We previously demonstrated that reducing Copia levels at the NMJ increases bouton formation, indicating that *Copia* acts as an inhibitor of synaptic plasticity (M’Angale et al., 2025). *Copia* protein levels are elevated in *Imp* mutant embryos and at the larval NMJ following *Imp* knockdown. Consistent with this upregulation, NMJs with reduced *Imp* levels exhibit fewer boutons compared to wild-type NMJs. Our findings suggest that loss of Imp may impair NMJ plasticity and potentially affect embryo survival through its regulatory role in controlling the expression of Copia and other transposons. It remains to be determined whether reducing *Copia* expression or the expression of other misregulated transposons can rescue the NMJ phenotype or mitigate age-related defects caused by *Imp* loss. We also found that *Profilin* mRNA, a regulator of the actin cytoskeleton and a previously described Imp target, is enriched in Imp-RNP complexes in embryos. In the brain, overexpression of *Profilin* mRNA can rescue the axonal remodeling defects observed in *Imp* mutants (Medioni et al., 2014). Whether *Profilin* mRNA influences axonal growth at the NMJ or interacts with *Copia* remains unclear but represents a highly interesting direction for future investigations.

The intricate regulation of TEs encompasses various mechanisms crucial for controlling their activity and preventing genomic instability. The piRNA (Piwi-interacting RNA) pathway is a primary and conserved mechanism governing TE expression in metazoans. In the cytoplasm, Piwi proteins and piRNAs cleave TE transcripts, suppressing their further transcription, expression, and transposition. The piRNA/Piwi machinery also operates within the nucleus, inducing histone H3 lysine 9 trimethylation (H3K9me3), ultimately driving transcriptional silencing of TEs. Several key players in the piRNA pathway—such as Piwi, Spn-E, Tdrd3, Hop, Aub, Ago3, and Cbp80—were enriched in the Imp-RNP complexes. Loss of *Imp* did not impair nuclear RNA export of piRNA precursors in oogenesis, and small RNA sequencing indicated that piRNA levels were not significantly altered in *Imp* mutant embryos, suggesting that *Imp*’s role in TE silencing might involve mechanisms independent of, or downstream from piRNA synthesis. However, we observed a strong effect on Piwi nuclear localization in the nurse cells of *Imp* mutant egg chambers, and Piwi protein was enriched in Imp IPs. Though the strong effect on Piwi localization observed in the absence of *Imp* in the germline contrasts with the lack of strong effects on Piwi localization during embryogenesis, zygotic transcription might mask the role of maternal Imp. Our results suggest that *Imp* may influence transposon silencing independently of piRNA production, potentially by ensuring the proper maternal deposition of Piwi or functional Piwi/piRNA complexes into early eggs, a hypothesis that requires further investigation.

The Imp interactions also point to another pathway. A complex comprising *Drosophila* Top3B, Tdrd3, and the RNAi-induced silencing complex (RISC)—housing Ago2, the p68 RNA helicase, and Fmr1—selectively associates with specific nascent RNAs featuring intricate tertiary structures. This complex contributes to resolving RNA topological challenges and recruits the heterochromatin components Su(var)3-9 and Su(var)205/HP1 (Lee et al., 2018). Together, they recruit methyltransferases pivotal for heterochromatin formation and transposon silencing. Interestingly, Fmr1, Top3B, Tdrd3, Su(Var)205/HP1, Aub, and His2A were associated with Imp in RNase-treated extracts, indicating direct binding of Imp to components of this transposon silencing complex, regardless of shared RNA or TE mRNA targets.

PSI (P-element somatic inhibitor), Hp48 (also known as Hrb27C), Hrp36 (Hrb87F), Hrp38 (Hrb98DE), and PABP were also enriched in Imp complexes. The presence of PSI in Imp complexes was specific, as point mutations in Imp’s dual KH domains, which lead to lethality similar to *Imp^2^*, *Imp^7^*, and *Imp^8^* mutants, also disrupted PSI binding (Fig. 8B). This complex functionally assembles on P-element pre-mRNAs, blocking splicing of the third intron (IVS3), preventing the production of fully spliced transposase mRNA, and the translation into an active transposase (Ghanim et al., 2020; Horan et al., 2015; Siebel et al., 1994; Siebel et al., 1995). This mechanism prevents transposition of P-elements in somatic cells. Recent research has shown that piRNAs regulate P elements and transposable elements, particularly the Gypsy retrovirus-like element, by controlling transposon alternative splicing (Ghanim et al., 2020; Karam Teixeira et al., 2017). This suggests that splicing regulation between P-elements and transposons may share similar mechanisms. An intriguing hypothesis is that Imp/PSI complexes may regulate transposon levels by controlling TE splicing. Moreover, Piwi protein and PSI levels were elevated in *Imp* mutant embryos (Table 1), suggesting that these embryos upregulate mechanisms to inhibit TE expression and to counteract the misexpression of genes involved in growth and development as an essential compensatory mechanism for embryo survival. Thus, *Imp* seems to impact embryo survival by affecting various RNA targets and controlling different RNA metabolic pathways.

The question of why Imp’s role in localizing certain developmental mRNAs appears redundant while its role in TE silencing is essential is intriguing. One possibility is functional compensation. Other RNA-binding proteins may be able to substitute for Imp in the BicD/Egl transport machinery for a subset of cargoes, ensuring robust developmental progression. However, its role in the piRNA pathway may be more specialized, perhaps involving specific protein-protein interactions or recognition of TE-associated RNA features that other proteins cannot replicate. This would make Imp an indispensable node in the TE silencing network, while being a more dispensable component of a broader RNA transport system.

A central finding of this study is the strong correlation between the loss of maternal Imp, the de-repression of TEs, and severe embryonic defects. While we propose that the uncontrolled activity of TEs is a primary driver of the developmental failure, we cannot formally exclude the possibility that TE de-repression is a secondary consequence of general cellular stress in a dying embryo. Disentangling cause and effect is a critical next step. A definitive experiment would be to perform a genetic rescue. If the developmental phenotypes of *Imp* mutant embryos could be partially or fully rescued by simultaneously silencing the most highly de-repressed transposons (e.g., using a multi-target RNAi approach), it would provide powerful evidence that TE activity is indeed the primary cause of lethality.

Our findings underscore the essential role of maternal Imp in early *Drosophila* development, likely through its roles in RNP complexes that regulate cell growth and differentiation, and by controlling transposon silencing. These processes contribute to a deeper understanding of the molecular pathways that govern embryogenesis and maintain genomic stability.

## METHODS

### Drosophila stocks

FRT lines *Imp^2^, Imp^8^*, and the original *Imp^7^* stock were kindly provided by Daniel S. Johnston (Munro et al., 2006). These lines were sequenced to confirm the nature of the mutations. *Imp^2^* carries a deletion of a region spanning the two putative AUGs, along with a 20Kb intronic insertion after the Imp start codons, which makes this line lethal. Although the deletion was as previously described, the additional insertion had not been reported before. *Imp^8^* contains a 6Kb insertion and a 16 bp short insertion, as well as a deletion of a region that removes the first KH domain. This contrasts with previously reported data, where the entire ORF was described as deleted. This line is also lethal. The *Imp^7^* line carries a deletion that starts after the two putative AUGs and extends through the rest of the ORF, deleting all KH domains and the PLD. This deletion extends further into the 3′ UTR as previously described, removing part of the 3′ UTR. A schematic representation of the mutant genes is shown in Fig. S5A. *Imp* germline clones were induced using the FLP/FRT OvoD technique. The *Imp^7^* line was recombined with the FRT chromosome *y,w,v,P{FRT(w{hs})}101* obtained from Bloomington (BL1844). Germline clones were then generated by crossing females bearing the *Imp^2^, Imp^8^* or *Imp^7^* FRT recombined chromosome *to w,ovoD1,v,P{FRT(w[hs])}101/C(1)DX,y,f/Y; P{hsFLP}38* males (Bloomington, BL1813). The progeny was heat-shocked twice a day for two days upon reaching the late L2 or early L3 stages. Females bearing both the *Imp* mutant chromosome and the *OvoD* chromosome were then used to collect embryos or ovaries lacking *Imp*.

*Imp-GFP^G080^* was kindly provided by William Chia and Xavier Morin (Morin et al., 2021). *Piwi^2^* and variegating white reporters P[w^var^], embedded in the fourth chromosome pericentric region (stock: *yw; P{hsp26-pt-T}* at *118E-10*, BL84108), the fourth chromosome heterochromatin (*P{hsp26-pt-T}39C-12 at 102B5*, BL84110), or the X chromosome (+w^m4h^, BL76618), were obtained from Bloomington. *Imp^H44^* and *Imp^H149^* flies were generously provided by T. Hays (Boylan et al., 2008). Cas9 fly stocks were obtained from Bloomington (BL78782 and BL54591). Transgenic flies were generated using the attP/attB system.(Bischof et al., 2007)

### Adult Drosophila aging assay

Adult fly aging was assayed with the previously described survival analysis (M’Angale and Staveley, 2016). In brief, single-vial matings with virgin females and males of the appropriate genotypes were performed at room temperature (RT) in fly rearing bottles. Age-matched cohorts of at least 200 F1 male flies were collected. The flies were pushed to new food every other day and scored for survival, with flies determined dead when they did not display movement upon agitation. Survival curves were graphed using GraphPad Prism software, and the curves were compared using the log-rank (Mantel-Cox) test.

### DNA constructs for RNA injections

DNA constructs used as templates to prepare RNA for injection experiments *hairy (h^SL1x3^), ftz* and *grk* were previously described (Bullock and Ish-Horowicz, 2001; Bullock et al., 2006) To generate the template to transcribe the *hairy (h^SL1x3^)* RNA containing a poly-(A)_71_ tail, a BamHI/HindIII fragment from the *h^SL1x3^* cDNA was subcloned into the NcoI/ BglII site of the FLuc-cassette plasmid (Hernández et al., 2004) by fill-in reaction to generate the *- h^SL1x3^ -(A)_71_* cDNA.

### DNA constructs for two-hybrid screens

The plasmids BicD carboxy-terminal (534-782)-BD, BicD full-length-AD, Egl full-length-AD and Egl full-length-BD were described (Vazquez-Pianzola et al., 2022). DNA fragments encoding *D. melanogaster* IMP were cloned in-frame with the Gal4 cDNA into the vector pOAD (activator domain, AD) to create the “bait” constructs (Cagney et al., 2000). Full-length *IMP1* was cloned as a EcoRI-NcoI fragment to create the construct IMP1 FL-AD. IMP1 amino-terminal (amino acids 1-234) was cloned as a EcoRI-PstI fragment to create the construct IMP1 (1-234)-AD. IMP2 amino-terminal (amino acids 1-242) was cloned as a EcoRI-PstI fragment to create the construct IMP2 (1-242)-AD, and IMP1 and IMP2 carboxy-terminal (amino acids 242-573 of IMP1, amino acids 235-566 of IMP2) were cloned as a EcoRI-NcoI to create the construct IMP1/2 (carboxy)-AD

### DNA constructs for generating the *Imp-GFP^Stop^* and *Imp^KH^* mutants

DNA constructs to generate the *Imp-GFP* plasmids for making the mutations in different KH domains were constructed as follows. The *Imp-GFP* region was amplified by PCR using genomic DNA obtained from *Imp-GFP* expressing flies and primers containing EcoRI and XhoI restriction sites (called IMP-rec-F and IMP-rec-R). The resulting fragments were cloned into the corresponding sites in pBluescript (pBSK). Primer sequences are listed in Table S9. The KH domains in the generated plasmid were then mutated by replacing the conserved GXXG motifs with GDDG sequences. These mutant protein versions have been shown to fold properly but impair *Imp* ortholog RNA binding (Hollingworth et al., 2012). These mutations were introduced either as single KH mutants or as double mutations in KH1 and KH2 (KH1-2), or in KH3 and KH4 (KH3-4), using site-directed mutagenesis with the corresponding primers listed in Table S9. Double mutants were generated by sequential mutations. The PAM sites corresponding to the gRNAs were also mutated in the donor DNAs.

Genomic sequences were scanned for suitable sgRNA target sites using FlyBase Jbrowse DNA oligos for cloning of the *gRNA* genes were ordered from Microsynth (Balgach, Switzerland) and cloned into pCFD5 following the protocols on the website: http://crisprflydesign.org/. Table S9 shows the primers used for cloning the different *gRNA* genes used in this study. gRNA-1 was used to generate the Imp-GFP^Stop^ mutant stock. Mid2 gRNA was used to generate the single KH mutations and the Mid1 gRNA for the generation of KH1-2 and KH3-4 double mutants.

### Generation of *Imp-GFP^Stop^* and *Imp^KH^* mutant flies

The *Imp-GFP^G080^* trap chromosome encodes a fusion of *Imp* with *GFP*, where the *GFP* is inserted in-frame after the 32nd amino acid of the Imp-RA isoform (Fig. S2) (Morin et al., 2001). Other isoforms could also be translated, differing only in the sequence of this short N-terminal region. After the GFP insertion site, both isoforms share the same sequences, including all the functional regions of the protein: the four KH domains and the conserved PLD domain. Moreover, homozygous *Imp-GFP^G080^* flies are viable and fertile (Boylan et al., 2008; Munro et al., 2006). The gRNA transgenes are described in the cloning section of the methods, and the sequences are shown in Table S9.

To generate a mutant version of the *Imp-GFP^G080^* chromosome, we performed a CRISPR-Cas9 mutagenesis screen using a gRNA targeting a site downstream of the GFP insertion in the Imp coding region (gRNA-1). The nos-Cas9 stock (BL54591) was used. This resulted in a mutant version of the *Imp-GFP* chromosome with an 8-nucleotide deletion (sequence: GACAGGGC), leading to a frameshift in the open reading frame and the introduction of a premature stop codon. The frameshift occurs after the 17th amino acid of the KH1 domain. This mutation was lethal in flies and resulted in the production of a GFP protein under the same regulatory elements as Imp expression. This fly line, referred to as *Imp-GFP^Stop^*, was used as a mock control for protein and RNA immunoprecipitations (Fig. S2A-B).

To generate *Imp-GFP^KH^* mutants in single KH domains, females bearing the Imp-GFP chromosome and the gRNA *Mid2* transgene were crossed with *nos-Cas9*-bearing males (BL54591). For the generation of double KH mutant stocks, females carrying the *Mid1,* gRNAs were crossed to males bearing the *Cas9* from the BL 78782 stock. The corresponding Imp-GFP mutated plasmids were injected into their embryo progeny. Single flies were then screened for the presence of the required mutations. Flies with single mutations, referred to as *Imp^KH1^*, *Imp^KH2^*, *Imp^KH3^*, and *Imp^KH4^*, as well as double mutant flies *Imp^KH1-2^* and *Imp^KH3-4^*, were generated by injecting the corresponding donor DNA *Imp-GFP* mutated templates. The donor plasmids were mutated in the corresponding KH domain as well as in the PAM site recognized by the gRNA used. Single mutant flies turned out to be viable and fertile, while flies bearing the double mutations were lethal.

### Yeast two-hybrid assays

Interactions between prey and bait proteins were tested in a yeast two-hybrid system using the haploid strains PJ69-4a and PJ69-4alpha (Cagney et al., 2000). Diploid cells containing both bait and prey plasmids were grown on selective media (―W(tryptophan), ―L (leucine)) and are shown as growth controls. Protein interactions were detected by replica-plating diploid cells onto the selective media (―W, ―L, ―H (histidine) + 5 mM 3-amino-1,2,4-triazole (3AT)). Growth was scored after 6 days of incubation at 30 °C.

### Protein Immunoprecipitations (IPs) and Western blot analysis

The immunoprecipitations (IPs), followed by mass spectrometry and Western blotting used to identify Imp in the BicD complexes, (depicted in Fig.s 1A-B) were previously partially described (Vazquez-Pianzola et al., 2011). Additionally, to detect Imp in the immunoprecipitates, anti-IMP antibodies (1:3,000, (Munro et al., 2006)) were used.

To identify novel Imp interactors, the IPs were performed essentially as described in (Vazquez-Pianzola et al., 2011) with some modifications. *Imp-GFP* and *Imp-GFP^Stop^* 0-8 h-old embryos were used. Dechorionated 0-8 h-old embryos were homogenized on ice in homogenization buffer (HB; 25 mM Hepes pH 7.4, 50 mM KCl, 1 mM MgCl2, 1 mM DTT, 0.5% NP-40, and EDTA-free protease inhibitor cocktail Complete™ (Roche Diagnostics)). 2 mL of HB was used per gram of embryos. The supernatants were centrifuged at 16,000 x g for 45 minutes at 4°C, and the resulting supernatant was transferred and centrifuged again at 16,000 x g for 25 minutes at 4°C. 2 mL of anti-GFP IDJ42 monoclonal antibody supernatant were incubated with 62.5 μL of protein-G Mag Sepharose magnetic beads (Cytiva) for 3 hours at room temperature. After extensive washing with PBS, the bead-antibody complexes were incubated overnight at 4°C on a rotating wheel with 1.7 mL of embryo extract per IP. The unbound material was washed off 7 times with 1 mL of homogenization buffer (HB) without detergents, and the beads were transferred to new tubes during the first and last washes. For Western blot analysis, a 100 μL aliquot of the resuspended beads was taken after the final wash and resuspended in 40 μL of sample buffer to confirm successful Imp pull-down. The remaining IP material was subjected to mass spectrometry analysis.

For Western blot analysis of Imp-GFP complexes, small-scale IPs were performed using 30-40 μL of beads and 1 mL of embryo extract. Blots were performed using the following primary antibodies: rabbit anti-Pabp (1:5,000 dilution), anti-Imp (1:3,000, (Munro et al., 2006), Fig. 1), anti-Imp (1:1,000,(Geng and Macdonald, 2006); Fig. 8), and mouse anti-BicD (a mix of 1B11 and 4C2, 1:10 dilution).

To detect Copia isoforms in Imp embryos, dechorionated embryos were homogenized in loading buffer containing 0.1 M DTT and 2% SDS, and the Copia isoforms were detected using anti-Copia antibodies (1:1,000, (M’Angale et al., 2025)). Horseradish peroxidase-conjugated antibodies were obtained from GE Healthcare.

### Whole embryo and IP mass spectrometry

For the preparation of whole embryo extracts for mass spectrometry analysis, 0-4 h-old dechorionated embryos were homogenized with a pestle in a buffer containing 8M Urea and 100 mM Tris-HCl (pH 8). The extract was centrifuged at 16,000 x g for 40 minutes at 4°C. Protein concentration was measured using the Bradford assay (BioRad). The supernatant was then subjected to mass spectrometry analysis. For the mass spectrometry identification of Imp interactors, IP beads were also subjected to mass spectrometry analysis. Mass spectrometry analysis was performed by the Proteomics and Mass Spectrometry Core Facility of the University of Bern. The proteins bound to the magnetic beads were resuspended in 8M Urea / 50 mM Tris-HCl (pH8), reduced for 30 min at 37^0^C with 0.1M DTT / 100 mM Tris-HCl pH8, alkylated for 30 min at 37^0^C in the dark with IAA 0.5M / 100 mM Tris-HCl (pH8) and diluted with 4 volumes of 20 mM Tris-HCl pH8 / 2 mM CaCl2 before overnight digestion at room temperature with 100 ng sequencing grade trypsin (Promega). Samples were centrifuged, and a magnet holder trapped the magnetic beads to extract the peptides in the supernatant. The digested proteins were analyzed by liquid chromatography. (LC)-MS/MS (PROXEON coupled to a QExactive mass spectrometer, ThermoFisher Scientific) with three injections of 5 μl digested proteins. Peptides were trapped on a μPrecolumn C18 PepMap100 (5 μm, 100 Å pore size, 300 μm x 5 mm, ThermoFisher Scientific, Reinach, Switzerland) and separated by backflush onto an analytical C18 column (5 μm, 100 Å, 75 μm x 15 cm, C18). Separation was performed using a 60 min gradient from 5% acetonitrile to 40% in water, 0.1% formic acid, at a flow rate of 350 ml/min. The Full Scan MS was acquired at a resolution of 70,000 with an automatic gain control (AGC) target of 1x10^6^ and a maximum ion injection time of 50 ms. The data-dependent method for precursor ion fragmentation was applied with the following settings: resolution 17,500, AGC of 1E05, maximum ion time of 110 milliseconds, mass window 2 m/z, collision energy 27, underfill ratio 1%, charge exclusion of unassigned and 1+ ions, and peptide match preferred, respectively. The data were then processed with the software MaxQuant (Cox and Mann, 2008), version 1.6.14.0, against the UniProtKB (Bateman, 2019)Drosophila 7228 database (release 2021_02) containing canonical and isoform entries, to which common contaminants were added. The following parameters were set: digestion by strict trypsin (maximum three missed cleavages), first search peptide tolerance of 15 ppm, MS/MS match tolerance of 20 ppm, PSM and protein FDR set to 0.01, and a minimum of 2 peptides requested per group. Carbamidomethylation on cysteine was selected as a fixed modification; the following variable modifications were allowed: methionine oxidation, deamidation of asparagines and glutamines, and protein N-terminal acetylation; a maximum of 3 modifications per peptide was allowed. Match between runs was turned on (match time windows 0.7 min) but only allowed within replicates of the same kind. Peptides were normalized by variance stabilization (Huber et al., 2002), imputed, and combined to form Top3 intensities (Silva et al., 2006) and considered alongside MaxQuant’s Label-Free Quantification (LFQ) values. Missing peptides, respectively LFQ intensities, were imputed by drawing values from a Gaussian distribution of width 0.3 centered at the sample distribution mean minus 2.5x the sample standard deviation, provided there were at most 1 non-zero value in the group; otherwise, the Maximum Likelihood Estimation (Silver et al., 2009) was used. Differential expression was performed by applying the empirical Bayes test (Kammers et al., 2015) between groups; significance testing was performed as described (Uldry et al., 2022).

### Immunostaining and RNA *in-situ* hybridization in embryos and ovaries

Immunostaining experiments were performed as described in previous studies (Rashpa et al., 2017; Vazquez-Pianzola et al., 2014; Vazquez-Pianzola et al., 2022). Immunostaining of whole-mount embryos and ovaries was conducted using the following primary antibodies: mouse anti-BicD (a mix of 1B11 and 4C2, 1:10 dilution), rabbit anti-Imp (1:1,500, (Geng and Macdonald, 2006)), rabbit anti-Imp (1:500, (Munro et al., 2006)), rabbit anti-Copia (1:500 (M’Angale et al., 2025)), mouse anti-Piwi P3G11 (1:1,000; (Saito et al., 2006)), mouse anti-Ago1 (1:500, (Miyoshi et al., 2005)), mouse anti-Hp1 (1:20, C1A9-S, Developmental Studies Hybridoma Bank), anti-H3K9m3 (1:600, AbCam 176916), anti-lamin ADL84 (1:500–1:300; Developmental Studies Hybridoma Bank), and anti-Serp (1:1,000; (Luschnig et al., 2006)). Secondary antibodies were obtained from Jackson ImmunoResearch or Molecular Probes. Where necessary, nuclei were stained with 2.5 μg/mL Hoechst 33258 (Molecular Probes) for 20 minutes during the final washing steps. Images were analyzed using a Leica TCS-SP8 confocal microscope. RNA *in situ* hybridization using Stellaris probes to detect piRNA precursors was performed essentially as described (Mohn et al., 2014; Rashpa et al., 2017). However, following hybridization, samples were subjected to immunostaining to reveal Imp and Lamin proteins. Stellaris probes for detecting the 42AB sense transcripts (probe 42AB-RS labeled with CalFluor 590) and probes against the 20A transcripts (labeled with Quasar 670) were also previously described in (Mohn et al., 2014). *In situ* hybridization experiments to detect mRNA expression were performed as described (Vazquez-Pianzola et al., 2017). Linearized *Osi9*, containing the *Osi9* cDNA, was used as a template to generate digoxigenin-labeled RNA antisense probes. Images were analyzed using a Leica TCS-SP8 confocal microscope and processed with Leica software, Photoshop, and ImageJ.

### NMJ immunostaining and protein intensity quantification

*Drosophila* larvae were dissected in HL3 media and fixed in 4% paraformaldehyde. Washes and permeabilization were performed in 0.1M PBS supplemented with 0.2% Triton X-100. Larval fillets were incubated overnight in primary and secondary antibodies at 4°C followed by 2 hours at RT. After this, they were mounted. Laser scanning confocal microscopy and intensity measurements were as previously described (M’Angale et al., 2025).

### Fluorescent RNA synthesis and injections

RNA for injections was synthesized in the presence of either Alexa-488-UTP (Molecular Probes) or Cy3-UTP (NE Life Sciences) as described (Bullock et al., 2006). *hairy* RNA was synthesized using linearized *h^SL1x3^*cDNA in pBSK^(1)^ or *h-(A)_71_* cDNA, *ftz* using *ftz*-pCRIITOPO and *grk* using pBSK-*grk* as templates. For blocking experiments, affinity-purified anti-Imp antibody (Munro et al., 2006) and anti-GFP (Roche or AMS Biotechnology) were pre-injected at 1 μg μl^-1^, 10-12 min before injecting the fluorescently labeled RNAs. Fluorescently labeled RNAs were used at 0.4 μg μl^-1^. Fixing and immunofluorescent stainings of injected embryos were done as described (Bullock and Ish-Horowicz, 2001; Vazquez-Pianzola et al., 2017). Images were analyzed with a Leica TCS-SP2 confocal microscope. Localization was designated as strong (most of the RNA was in the apical cytoplasm), weak (most of the RNA remained basal, but there were some apical caps of RNA) or unlocalized (no apical concentration of RNA).

### Pigment determination

Pigment quantification was performed as described in (Rabinow et al., 1991). The heads of 22 flies of the appropriate genotypes were dissected and frozen on dry ice. The heads were homogenized with a pestle in 550 μL of methanol acidified with 0.1% HCl, followed by the addition of another 550 μL of acidified methanol. The extract was briefly centrifuged for 1 minute at 16,000 x g, and the supernatant was used to measure absorbance at 480 nm.

### Isolation of Imp-bound RNAs

RNA immunoprecipitation (RNA-IP) was essentially performed as described in (Vazquez-Pianzola et al., 2017) with the following modifications. Flies of the desired genotypes (*Imp-GFP^G80^, Imp-GFP^Stop^/Fm7c*) were reared in population cages. For each IP, 1.2 mL of 0-8 h-old embryo extract (dechorionated embryos homogenized at a ratio of 1 g embryos to 2 mL of homogenization buffer) of the corresponding genotype was mixed with 646 μL of adjusting buffer. The IPs were performed in triplicates, using embryos from different collections. For each IP, 80 μL of magnetic beads (Protein G Mag Sepharose, Cytiva) were used. A detailed protocol is available upon request.

### RNA purification, Library Preparation, RNA sequencing and analysis

Total RNA was isolated using TRIzol® Reagent (Invitrogen) following the manufacturer’s instructions. For transcriptome and small RNA analysis, 0-4 or 0-8 h-old embryos were collected, dechorionated, and homogenized directly in the indicated amount of TRIzol reagent. For RIP-Seq, TRIzol reagent was added directly to the RNA bound to the magnetic beads. RNA concentration was measured using a Qubit RNA BR or HS Assay Kit (Thermo Fisher Scientific, Q10211 or Q32855, respectively) in a Thermo Fisher Scientific Qubit 4.0 fluorometer. RNA quality was assessed using the Agilent Advanced Analytical Fragment Analyzer System with a Fragment Analyzer RNA Kit (Agilent, DNF-471).

For RIP-Seq and transcriptome analysis, library preparation and RNA sequencing were performed at the Next Generation Sequencing Platform of the Vetsuisse Faculty in Bern. Libraries for RNA-seq were prepared using Lexogen’s CORALL Total RNA-Seq Library Prep Kit with RiboCop (Lexogen, 117.96). The libraries were paired-end sequenced with a read length of 100 bp on an Illumina NovaSeq 6000 instrument (100 cycles; Illumina). The quality of the RNA-seq data was assessed using fastqc v.0.11.9 (http://www.bioinformatics.babraham.ac.uk/projects/fastqc/) and RSeQC v.4.0.0 (Wang et al., 2012). The reads were mapped to the reference genome using HiSat2 v.2.2.1 (Kim et al., 2015). FeatureCounts v.2.0.1 (Liao et al., 2014) was used to count the number of reads overlapping with each gene as specified in the genome annotation (Drosophila_melanogaster.BDGP6.32.103) The Bioconductor package DESeq2 v1.38.3 (Love et al., 2014) was used to test for differential gene expression between the experimental groups. ClusterProfiler v4.7.1 (Wu et al., 2021) was used to identify gene ontology terms containing unusually many differentially expressed genes. An interactive Shiny application v1.6.0 (Chang et al. 2022) was set up to facilitate the exploration and visualization of the RNA-seq results. All analyses were run in R version 4.2.1 (2022-06-23) (R Core Team 2022). The small RNA libraries were prepared and sequenced by a Novogene sequencing platform. Single-end 50 bp sequencing strategy was used to sequence the samples. The sRNAPipe in Galaxy was used for small RNA analysis (Pogorelcnik et al., 2018). The mapping was done, allowing either 1 or 2 mismatches. The small RNAs were mapped to the *Drosophila melanogaster* BDGP6.46 version.

## ACKNOWLEDGEMENTS

We thank Drs. Daniel St Johnston, Thomas Hays, Fabio Mohn, Mikiko C. Siomi, and Nicola Iovino for providing antibodies, probes, constructs, and fly stocks. We also thank Alexander Nater, Geert van Geest, and Rémy Bruggmann from the Interfaculty Bioinformatics Unit for their assistance with the RNA-seq differential expression analysis. Additionally, we express our gratitude to Dr. Pamela Nicholson from the NGS Platform for her advice on RNA-seq protocols. We acknowledge the Developmental Studies Hybridoma Bank (DSHB, University of Iowa) for antibodies, the Bloomington Stock Center (University of Indiana) for fly stocks, and FlyBase for invaluable support.

## SUPPLEMENTARY FIGURE LEGENDS

**Fig. S1.**
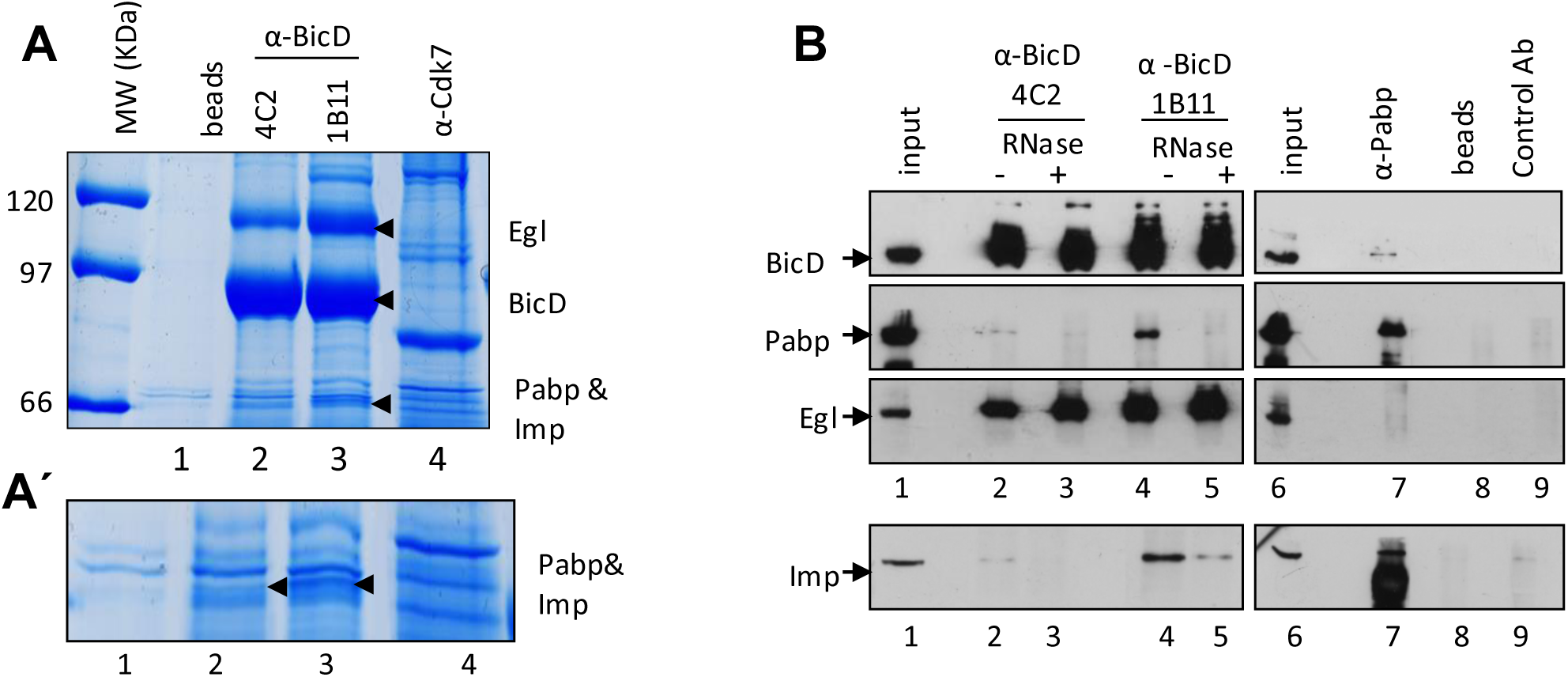
Imp forms part of the BicD/Egl complex and the interaction is stabilized by the presence of RNA. **(A-B)** Immunoprecipitation (IP) of total embryo extracts was done with the indicated antibodies. The IP material was resolved by SDS-PAGE and stained with Coomassie Blue **(A)** or analyzed by Western blot to detect the indicated proteins **(B)**. **(A)** Anti-BicD antibodies 4C2 (lane 2) and 1B11 (lane 3), which recognize different epitopes, were used for the IPs. Anti-Cdk7 antibody (unrelated control protein, lane 4) and protein G beads alone (lane 1) were used as controls for nonspecific binding. The gel area containing Pabp, Imp, BicD, and Egl is shown; the bands containing these proteins are indicated by arrowheads**. (Á)** Magnification of the gel region where Pabp and Imp were identified. **(B)** IPs performed with anti-BicD 4C2 (lanes 2 and 3), anti-BicD 1B11 (lanes 4 and 5), anti-Pabp (lane 7), beads alone (lane 8), and a control monoclonal antibody against a murine protein (lane 9). 0.15% of the cytoplasmic extract used for each IP was loaded as an input control (lanes 1 and 6). The IPs were performed in the presence (+, lanes 3 and 5) or absence (-, lanes 2, 4, 7, 8, and 9) of RNase A. These figures were partially published in (Vazquez-Pianzola et al., 2011) and are reproduced in this report since Imp was identified by mass spectrometry on the same band where Pabp was detected.

**Fig. S2.**
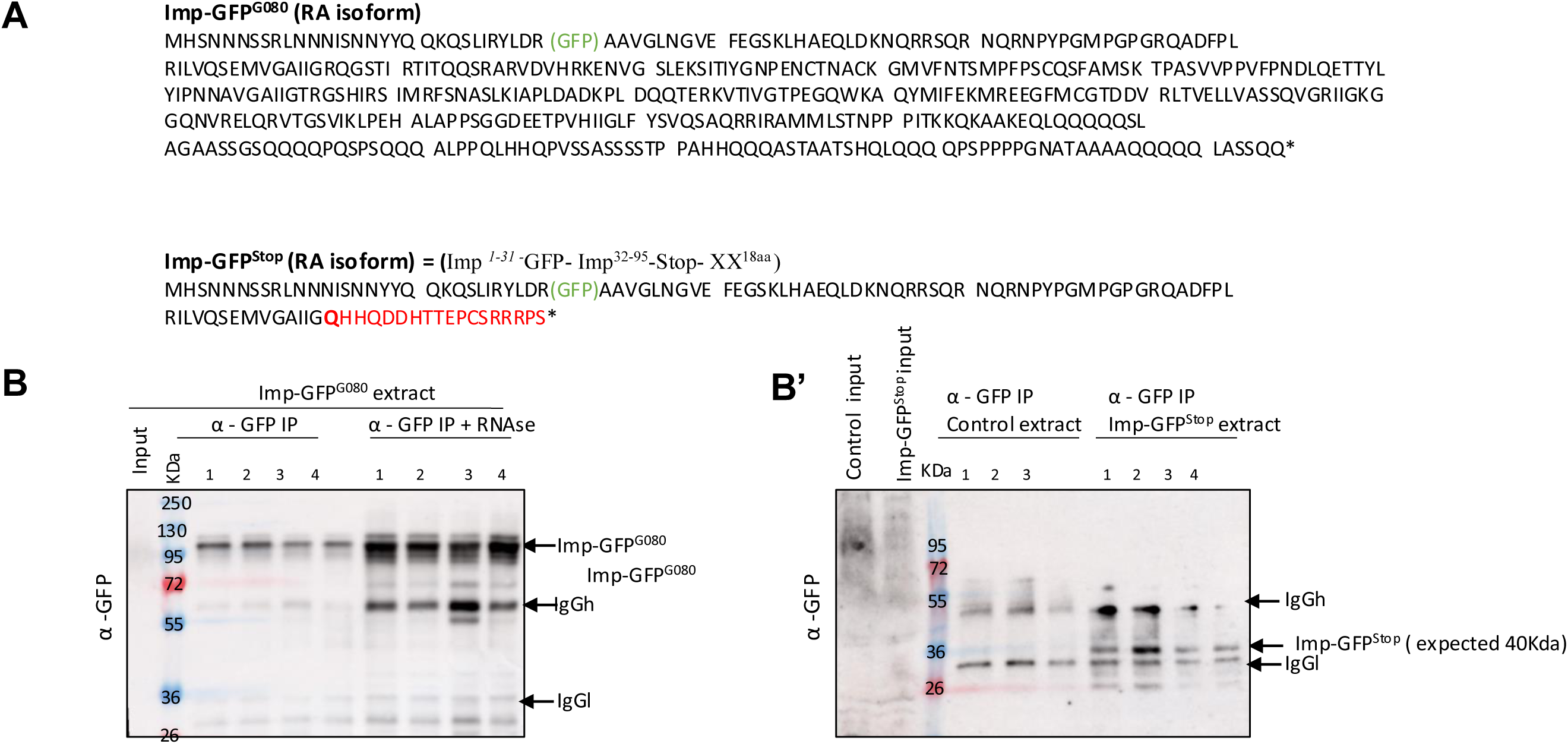
Structure and expression of the *Imp-GFP* genes. **(A**) Sequence of the RA protein isoform predicted to be generated from the *Imp-GFP^G080^* chromosome, and the mutant version produced via CRISPR-Cas9 mutagenesis of the same chromosome, referred to in the text as the *Imp-GFP^Stop^* line. Both alleles were sequenced. **(B-B’)** Input and IP samples were loaded, and Imp fusion proteins were detected using anti-GFP antibodies. Control extract, not expressing GFP was prepared from *yw* embryos. 1% of the input and 2.5% **(B)** and 6% **(B’)** of the IP material were loaded. The fusion proteins were not detected in the input samples due to their high dilution. However, specific bands corresponding to the expected molecular weight of the fusion proteins were detected in the IP samples. The heavy and light chains of the monoclonal antibody, also recognized by the secondary anti-mouse antibody, are indicated.

**Fig. S3.**
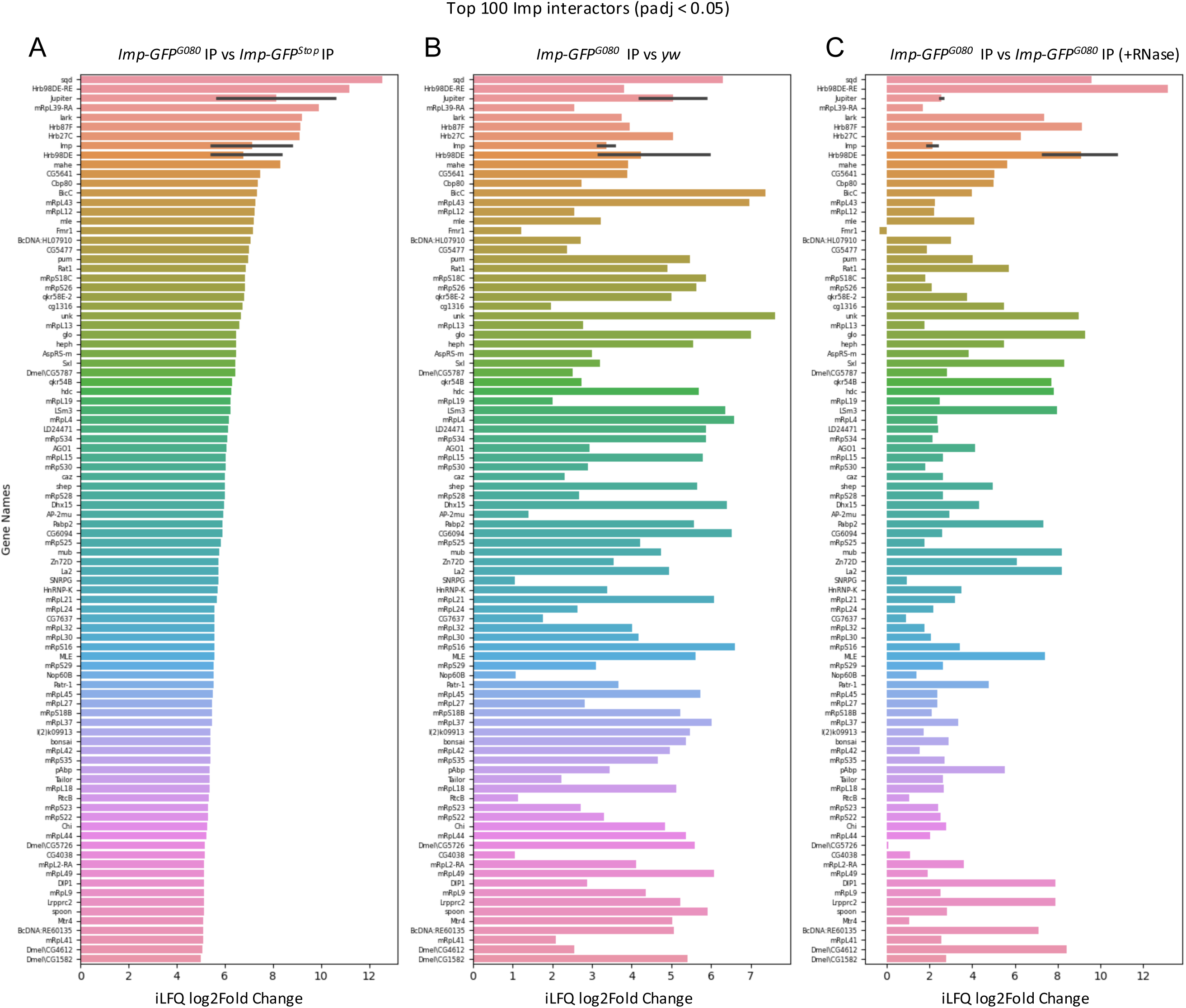
Identification of *Imp*-associated proteins. Embryo extracts from *Imp*-GFP*^G080^* (expressing full-length *Imp*-GFP) and control extracts used for mock IPs - Imp-GFP^Stop^ (expressing a truncated version of Imp internally tagged with GFP) and *yw* (not expressing any GFP) - were subjected to immunoprecipitation (IP) using anti-GFP antibodies. Additionally, *Imp*-GFP^G080^ extracts were subjected to IP with anti-GFP antibodies in the presence of RNase to detect RNA-independent interactors. The IP materials were directly analyzed by bulk mass spectrometry. **(A, B)** Top enriched proteins showing the highest log2FoldChange between the *Imp*-GFP pull-downs and the control mock pull-downs. **(C)** Top enriched proteins in the *Imp*-GFP pull-downs conducted in the absence vs. presence of RNase. A cutoff of padj < 0.05 was also applied. Pull-downs were performed in biological triplicates.

**Fig. S4.**
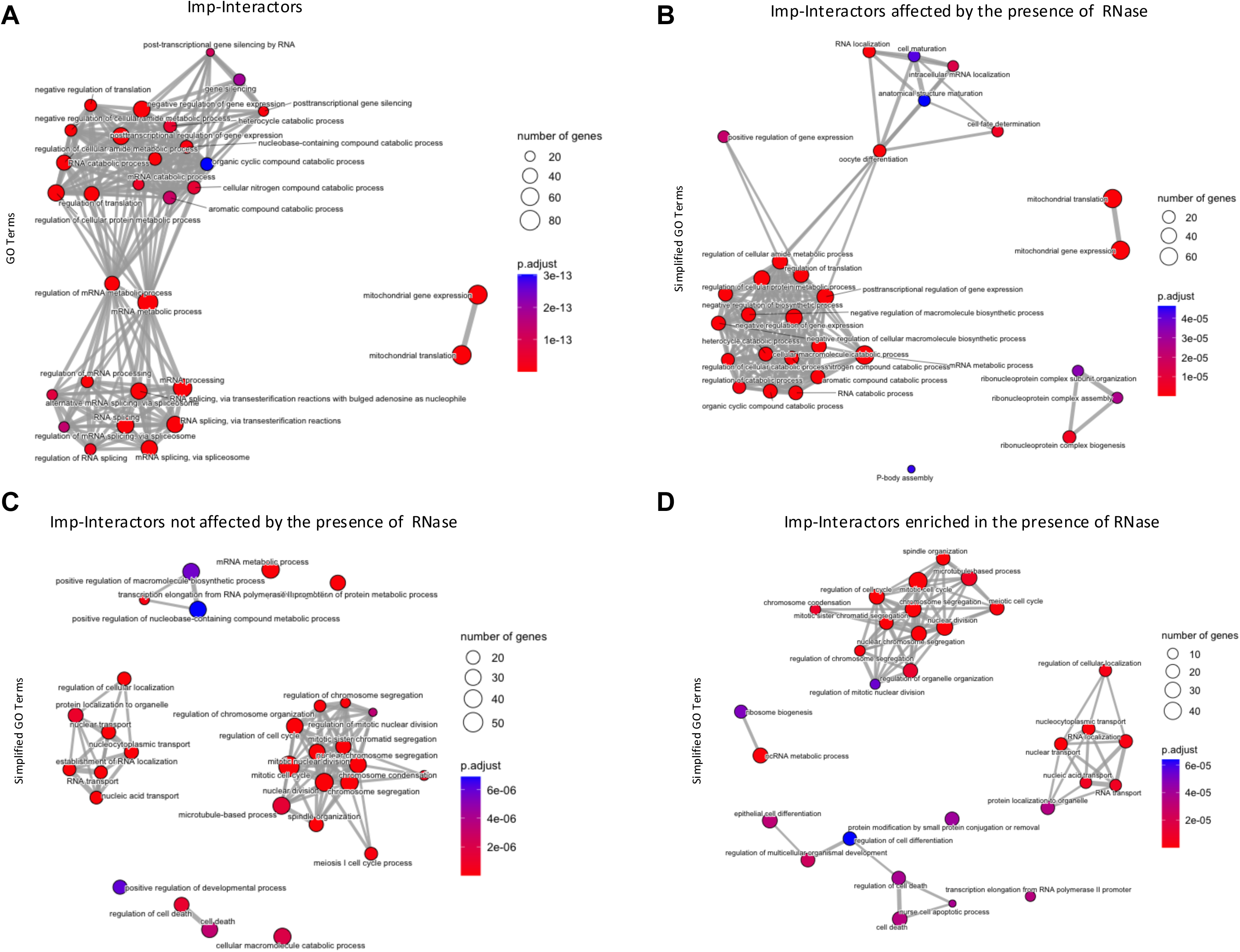
GO analysis of Imp interactors. Embryo extracts from *Imp-GFP^G080^*, *Imp-GFP^Stop^* or *yw* were subjected to immunoprecipitation (IP) using anti-GFP antibodies. Additionally, *Imp-GFP^G080^* extracts were subjected to IP with anti-GFP antibodies in the presence of RNase to detect direct interactors. The IP materials were directly analyzed by bulk mass spectrometry. The top 30 GO terms (A) and Simplified GO Terms (B-D), identify with Cluster profiler, are depicted. **(A)** GO analysis of the proteins enriched in the *Imp-GFP^G080^* IPs compared to the mock IPs (performed with *yw* and *Imp-GFP^Stop^* extracts). Interacting proteins were selected based on a log2Fold Change >1 or were uniquely present in the *Imp-GFP ^G080^* IPs, with a padj <0.05. **(B)** GO analysis of proteins associated with Imp via RNA (i.e., their interaction was lost in the presence of RNase). These proteins were preselected as in **(A)** but showed a reduced association with Imp in the presence of RNase. **(C)** GO analysis of proteins either enriched (log2Fold Change >1) in the *Imp-GFP^G080^* IPs performed in the presence of RNase (compared to the one performed in the absence of RNase) or that did not show changes (no significant padj value). The proteins are also enriched (log2Fold Change >1) compared to the mock *Imp-GFP^Stop^* IP (padj <0.05 for both). **(D)** GO analysis of proteins enriched (log2Fold Change >1) in the *Imp-GFP^G080^* IP performed in the presence of RNase compared to the one performed in the absence of RNase. These proteins are also enriched (log2Fold Change >1) compared to the mock *Imp-GFP^Stop^* IP (padj <0.05 for both).

**Fig. S5.**
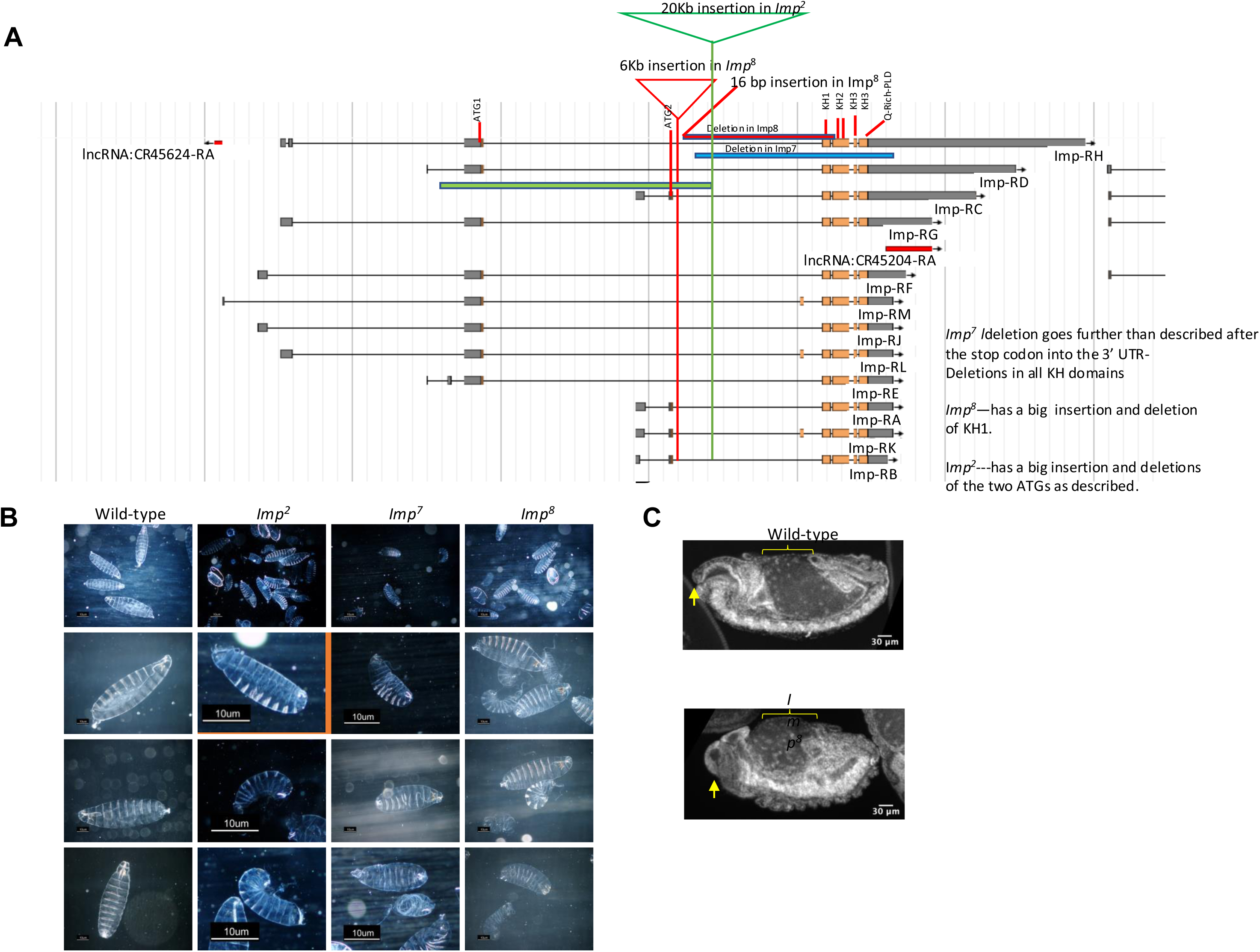
Embryos lacking maternal *Imp* arrest in early development, exhibiting anterior defects and defective dorsal closure. **(A)** The diagrams show the *Imp* gene structure and the alternatively spliced isoforms. Exons are depicted as boxes, introns as lines. Protein-coding exons are marked in orange. The deletions and insertions identified in the *Imp^2^*, *Imp^8^*, and *Imp^7^*alleles are depicted in green, red, and blue, respectively. **(B)** Cuticle preparations of embryos devoid of maternal *Imp* (derived from *Imp^2^, Imp^7^,* or *Imp^8^*germline clones) and *Imp^+^* controls (*yw*). **(C)** Embryos derived from *Imp^8^* GLCs and wild-type embryos were stained with Hoechst to visualize nuclear DNA to assess embryo morphology.

**Fig. S6.**
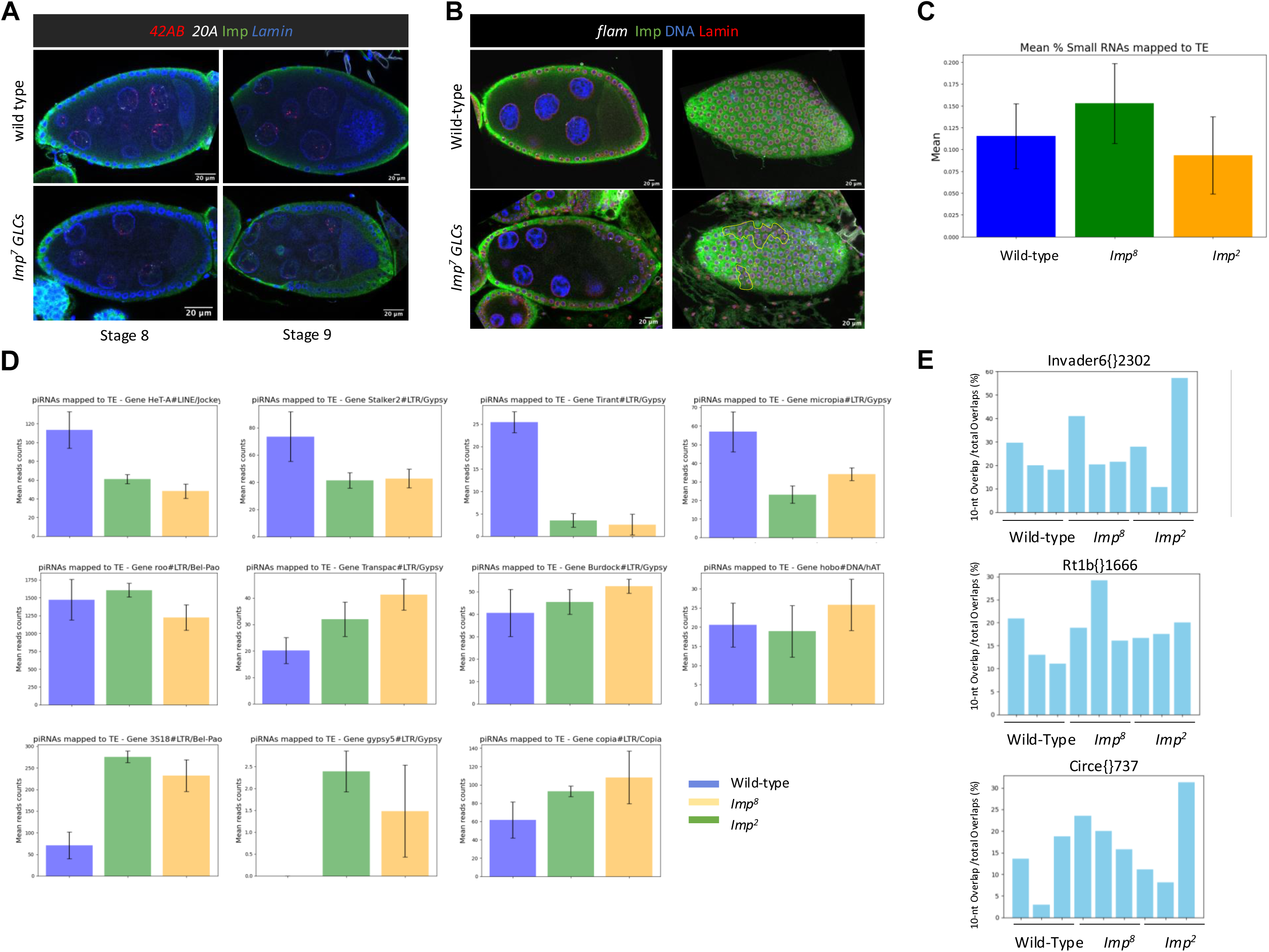
Nuclear RNA export of piRNA precursors and production of piRNAs mapping to deregulated TEs are not impaired in embryos devoid of *Imp*. **(A-B)** Wild-type and *Imp7* mutant ovaries generated by GLC induction were stained for Imp (green). **(A)** The ovarioles were additionally stained for Lamin (blue) and piRNA precursor transcripts from clusters 42AB (red) and 20A (white). Egg chamber stages 8 and 9 are shown for the different genotypes. **(B)** The ovaries were also stained for Lamin (red) and *flamenco* (*flam*) locus precursor RNAs. Egg chambers are shown in two different planes: one focusing on the nurse cells and oocyte to show the reduction of the Imp signal in the oocyte observed in clones devoid of Imp in the germline, and the other focusing on the follicle cells to show the presence of *Imp* mutant somatic clones marked within the yellow line. Although the precursor signals were not quantified, no visual differences were observed between the different genotypes. **(C)** Small RNAs purified from *Imp^+^* (*yw*) embryos and embryos lacking maternal Imp (by induction of *Imp8* and *Imp2* GLCs) were quantified by sequencing. The small RNAs were mapped to TE RNAs. The experiment was performed in biological triplicates. No significant difference in the amount of small RNAs mapping to TEs was observed between the mutants and the controls. The mean read counts mapping to TE sequences are shown for the different genotypes. **(D)** Mapping of piRNAs to specific TEs. The mean read counts of piRNAs mapping to different TE sequences are shown. **(E)** The Ping-Pong cycle is not clearly affected in *Imp* mutants since the percentage of sense and antisense piRNA pairs with a 10-nt overlap, relative to the total piRNA pairs, does not show a clear difference between *Imp^+^* and *Imp* mutant samples. Only piRNAs for which the sum of overlaps across all samples is 10 or more are shown.

**Fig. S7.**
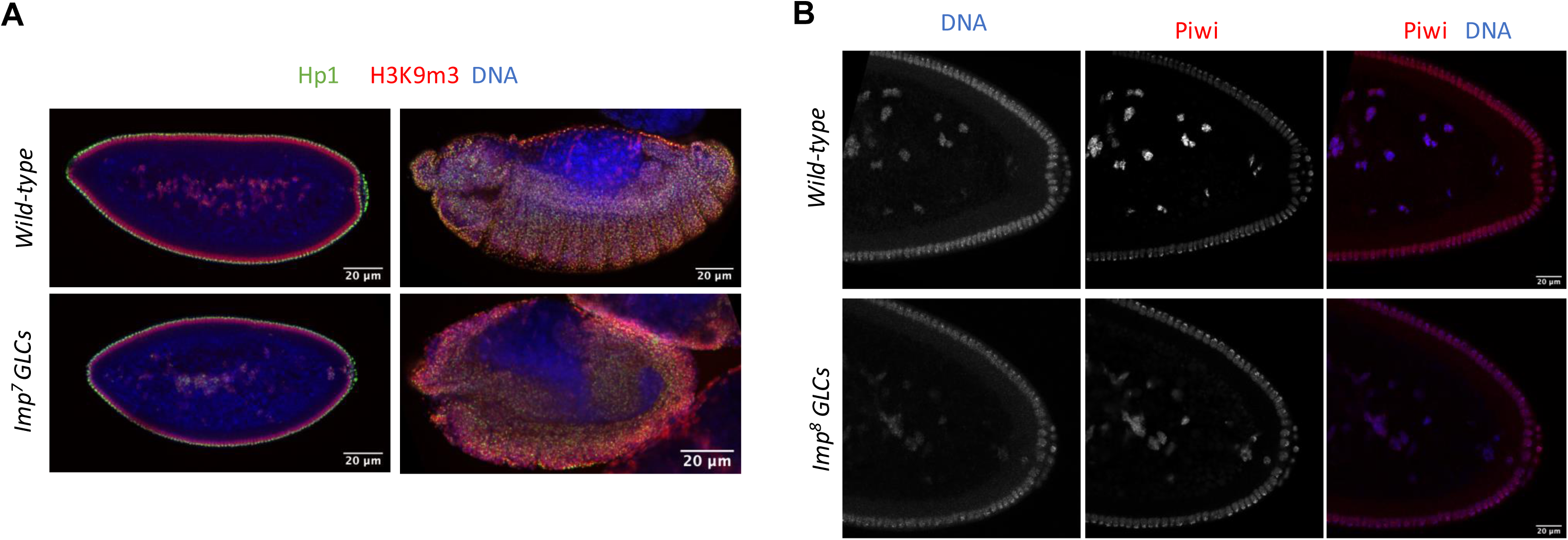
Embryos lacking maternal *Imp* do not show abnormal Piwi, Hp1, or H3 lysine 9 trimethylation staining signals. Maternal *Imp*-depleted and *Imp^+^*embryos were immunostained for Hp1 (green) and for histone H3 lysine 9 trimethylation (H3K9me3, red) **(A),** or for Piwi protein (red) **(B).** The embryos were also counterstained with Hoechst to visualize the DNA (blue).

**Fig. S8.**
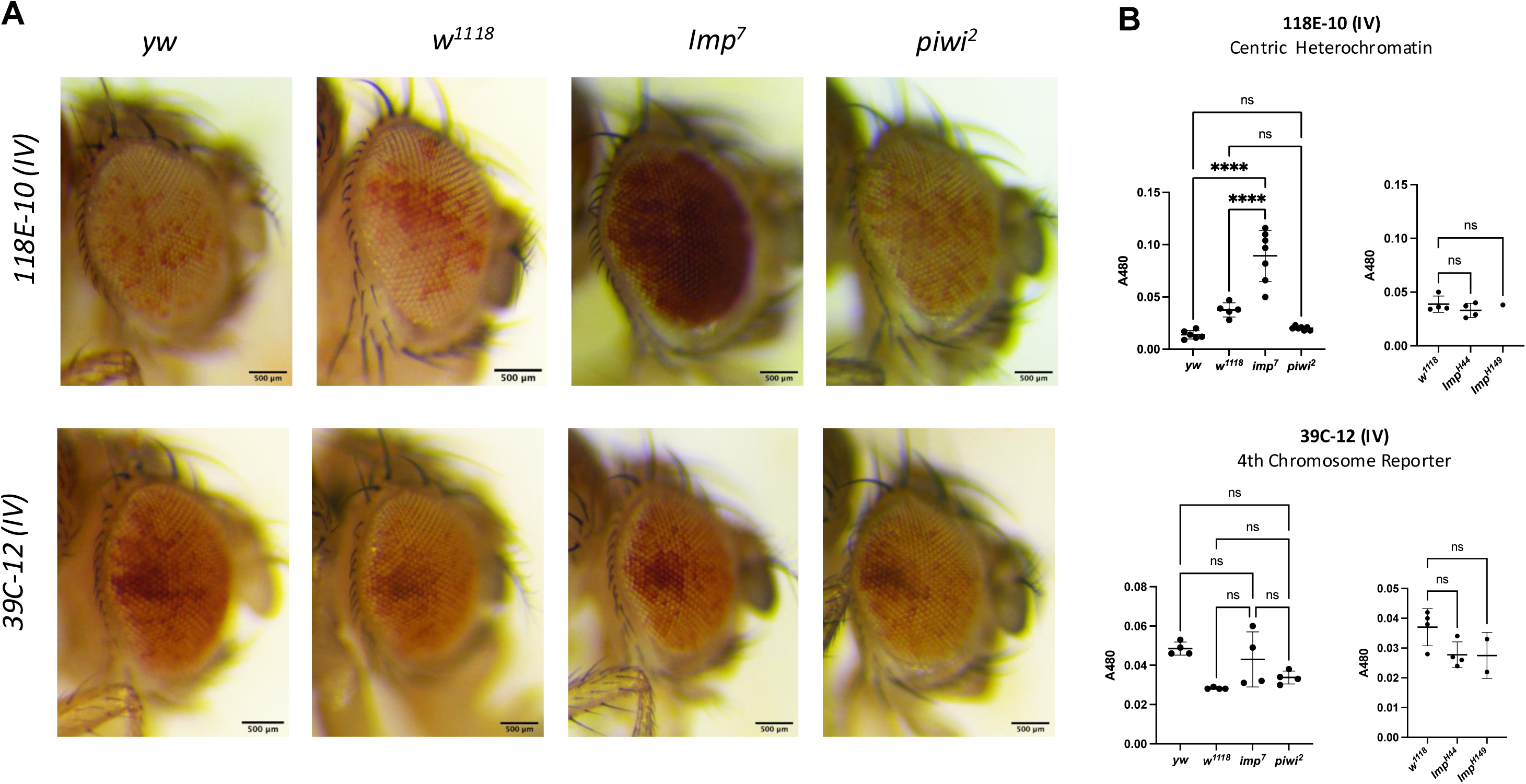
*Imp* loss in transcriptional silencing of different *white* reporters by position-effect variegation (PEV) **(A)** Representative images of eye pigmentation observed in the indicated genetic backgrounds. The 118E-10 (IV) *white* reporter line in the centric heterochromatin and the 39C-12 (IV) reporter in the 4th chromosome were used. The *Imp* mutant lines, *Imp^7^, Imp^H44^,* and *Imp^H14^*, as well as the *piwi* mutant, *piwi^2^*, were tested for their capacity to silence the variegating *white* gene. *Imp^H44^* and *Imp^H14^* are excision lines generated by the mobilization of a different P-element than the one used to generate the *Imp^7^* allele. **(B)** Eye pigment quantification for the indicated genotypes was measured by absorbance at A480 nm. None of the tested alleles showed an effect on the 4^th^ chromosome reporter expression. Although *Imp^7^* shows a significant de-repression of transcriptional silencing by PEV on the heterochromatin 118E-10 (IV) reporter compared to the *yw* and *w^118^*controls, neither the other *Imp* mutant alleles nor *piwi^2^*showed this difference. This suggests that the defective transcriptional silencing by PEV of this particular *white* reporter could be due to a different genetic background in the *Imp^7^*mutant.

