## Supplemental Table S9 for "Imp, a key regulator of transposable elements, cell growth, and differentiation genes during embryogenesis"

Supplementary Table S9. **List of primers used in this study and described in the Methods section.**

| **Primer Name** | **Sequence** | **Used for:** |
| --- | --- | --- |
| IMP-rec-F | 5'-TGG ATC CCC CGG GCT GCA GGA A TT CGG CAT ACC GAT GCA AAC AC-3' | Imp-GFP genomic region template used for homologous recombination-forward primer |
| IMP-rec-R | 5'-ATT GGG TAC CGG GCC CCC CCT CGA GTG GTT TCT ATG AAC T GAT AGC G-3' | Imp-GFP genomic region template used for homologous recombination- reverse primer |
| KH1 DD sense | 5'-GGCGCCATCATT**GGT**GATGAC**GGC**AGCACCATCAGG-3' | GRQG to GDDG  In KH1 domain |
| KH1 DD anti | 5'-CCTGATGGTGCTGCCGTCATCACCAATGATGGCGCC-3' | GRQG to GDDG  On Imp KH1 domain |
| KH2 DD sense | 5'-TGATTGGACGAATCATT**GGC**GATGAC**GGC**AATACCATTAAACGGATC  -3' | GKSG to GDDG mutation on Imp KH2 domain |
| KH2 DD anti | 5'-GATCCGTTTAATGGTATTGCCGTCATCGCCAATGATTCGTCCAATCA  -3' | GKSG to GDDG mutation on Imp KH2 domain |
| KH3 DD sense | 5'-GGCGCCATTATC**GGC**GACGAT**GGC**TCGCATATCCGAAG-3' | GTRG to GDDG mutation on Imp KH3 domain |
| KH3 DD anti | 5'-CTTCGGATATGCGAGCCATCGTCGCCGATAATGGCGCC-3' | GTRG to GDDG mutation on Imp KH3 domain |
| KH4 DD sense | 5'-GGGCCGTATCATT**GGC**GACGAT**GGC**CAAAATGTGCGAGA-3' | GKGG to GDDG mutation on Imp KH4 domain |
| KH4 DD anti | 5'-TCTCGCACATTTTGGCCATCGTCGCCAATGATACGGCCC-3' | GKGG to GDDG mutation on Imp KH4 domain |
| Mid2-PAM mut sense: | 5'-CATG TCG CGT GCT GAG AA**T** CAA ATT AGC ACC AAG CTG CG-3' | Mutation of PAM site in donor DNAs for Mid2 gRNA cutting. Blue=PAM site mutation, Red=Mid2 gRNA, sense primer |
| Mid2-PAM mut anti | 5'-CGCAGCTTGGTGCTAATTTGATTCTCAGCACGCGACATG-3' | Mutation of PAM site in donor DNAs for Mid2 gRNA cutting-antisense primer |
| Mid1-PAM mut fwd | 5'-CTC ATG CCC GGT CTC CAC CCT ATG GCA ATG ATG TCG ACA CC-3' | Mutation of PAM site in donor DNAs for Mid1 gRNA cutting-antisense primer |
| Mid1-PAM mut rev | 5'-GGT GTC GAC ATC ATT GCC ATA GGG TGG AGA CCG GGG AAC ATG AG-3’ | Mutation of PAM site in donor DNAs for Mid1 gRNA cutting-antisense primer |
| Imp-gRNA-1 fwd | 5'-TGC AGG GCG CCA TCA TTG GTC GAC-3' | For cloning the gRNA-1 used for generating Imp-GFP^stop^ stock |
| Imp-gRNA-1 rev | 5'-AAA CGT CGA CCA ATG ATG GCG CCC-3 | For cloning the gRNA-1 used for generating Imp-GFP^stop^ stock |
| Imp-gRNA-Mid2 fwd | 5'-TGC AGG CGC AGC TTG GTG CTA A TT-3' | For cloning the gRNA-Mid2 used for generating Imp-GFP^KH^ single domain mutants stocks |
| Imp-gRNA-Mid2 rev | 5'-AAA CAA TTA GCA CCA AGC TGC GCC-3' | For cloning the gRNA-Mid2 used for generating Imp-GFP^KH^ single domain mutants stocks |
| Imp-gRNA-Mid1 fwd | 5'-TGC AGT TCC CCG GTC TCC ATC CGA-3' | For cloning the gRNA-Mid1 used for generating Imp-GFP^KH^ double domain mutants stocks |
| Imp-gRNA-Mid1 rev | 5'-AAA CTC GGA TGG AGA CCG GGG AAC-3 | For cloning the gRNA-Mid1 used for generating Imp-GFP^KH^ double domain mutant stocks |
| IMP-EcoRI-FW | 5’-GAATTCATGCACAGCAACAATAATAG-3’ | Sense primer. For cloning the IMP-1 cDNA from the ATG used for the 2Hybrid screen. |
| IMP-1/2-NcoI-REV | 5’-CCATGGTTACTGTTGTGAGCTCGCCA-3’ | Reverse primer. For cloning the IMP-1 and 2 cDNAs, including the stop codon, common to both IMP isoforms. Nt 1701 (aa 566) of IMP-1; Nt 1722 (aa 573) of IMP-2. Used for the 2Hybrid screen. |
| IMP-2 EcoRI FW | 5’GAATTCATGGCATCCGAACTGGATCAATTC3’ | Sense primer. For cloning the IMP-2 cDNA from the ATG. Used for the 2Hybrid screen. |
| IMP-1/2 PstI REV | 5’-CTGCAGGTTCTCGTAGCTCTGGCGCAGCTTG-3’ | Reverse primer. For cloning the amino-terminal of both IMP-1 and 2 cDNAs , common to both IMP isoforms. Nt 702 (aa 234) of IMP-1; Nt IMP-1; Nt 723 (aa 241) of IMP-2. Used for the 2Hybrid screen. |
| IMP-1/2 EcoRI FW | 5’-GAATTCGATCTGCAGGCGATGGCGCCGCAGA-3’ | Sense primer. For cloning the amino-terminal of both IMP-1 and 2 cDNAs, common to both IMP isoforms. Nt 703 (aa 235) of IMP-1; Nt IMP-1; Nt 724 (aa 242) of IMP-2. Used for the 2Hybrid screen. |
